# A new mouse model of ATR-X syndrome carrying a common patient mutation exhibits neurological and morphological defects

**DOI:** 10.1101/2023.01.25.525394

**Authors:** Rebekah Tillotson, Keqin Yan, Julie Ruston, Taylor de Young, Alex Córdova, Valérie Turcotte- Cardin, Yohan Yee, Christine Taylor, Shagana Visuvanathan, Christian Babbs, Evgueni A Ivakine, John G Sled, Brian J Nieman, David J Picketts, Monica J Justice

## Abstract

ATRX is a chromatin remodelling ATPase that is involved in transcriptional regulation, DNA damage repair and heterochromatin maintenance. It has been widely studied for its role in ALT-positive cancers, but its role in neurological function remains elusive. Hypomorphic mutations in the X-linked ATRX gene cause a rare form of intellectual disability combined with alpha-thalassemia called ATR-X syndrome in hemizygous males. Patients also have facial dysmorphism, microcephaly, musculoskeletal defects and genital abnormalities. Since complete deletion of ATRX in mice results in early embryonic lethality, the field has largely relied on conditional knockout models to assess the role of ATRX in multiple tissues. Given that null alleles are not found in patients, a more patient-relevant model was needed. Here, we have produced and characterised the first patient mutation knock-in model of ATR-X syndrome, carrying the most common patient mutation, R246C. This is one of a cluster of missense mutations located in the chromatin interaction domain that disrupts its function. The knock-in mice recapitulate several aspects of the patient disorder, including craniofacial defects, microcephaly and impaired neurological function. They provide a powerful model for understanding the molecular mechanisms underlying ATR-X syndrome and for testing potential therapeutic strategies.

## Introduction

*ATRX* (alpha-thalassemia/intellectual disability, X-linked) was initially identified as a trans-acting factor which when mutated downregulates alpha-globin expression. Such mutations are responsible for a rare condition in boys defined by the combination of alpha-thalassemia and intellectual disability (ATR-X syndrome) (Gibbons et al., 1995). To date, ATRX has been reported to have roles in heterochromatin maintenance, epigenetic patterning, transcriptional regulation, and DNA damage repair (Timpano and Picketts, 2020). Since the discovery that ATRX is often lost in ALT- (alternative lengthening of telomeres) positive tumours (Heaphy et al., 2011), studies on the molecular mechanism by which ATRX supresses the ALT pathway in cancer (Kent and Clynes, 2021) far outnumber those on its roles in development.

ATR-X syndrome (Mendelian Inheritance in Man, 301040) is caused by hemizygous mutations in the X-linked *ATRX* gene in males. Mutations are often inherited from asymptomatic carrier females, who are protected by skewed X chromosome inactivation. All patients develop mild-severe intellectual disability and 75% of patients have alpha-thalassemia with variable degrees of severity. The condition is also characterised by morphological features, including facial dysmorphism, microcephaly, short stature, hypotonia, kyphosis, scoliosis and genital abnormalities (Gibbons, 2012; Stevenson, 2020).

Global deletion of *Atrx* in mice results in early lethality at E9.5, due to trophectoderm defects (Garrick et al., 2006). Therefore, the field has largely relied on conditional knock-out mice to investigate the role of ATRX in multiple tissues. Deletion of ATRX in the whole central nervous system or in the forebrain (by *Nestin-Cre* and *Foxg1-Cre*, respectively) result in postnatal lethality within 48 hours of birth (Bérubé et al., 2005). Other models are viable into adulthood, with phenotypes revealing the importance of ATRX for the proper development and function of neurons (Tamming et al., 2017; Tamming et al., 2020), the retina (Lagali et al., 2016; Medina et al., 2009), chondrocytes (Solomon et al., 2009), skeletal muscle (Huh et al., 2012; Huh et al., 2017), and Sertoli cells (Bagheri-fam et al., 2011).

The *ATRX* gene is comprised of 35 exons, spanning ∼280 kb, and encodes a protein 2,492 amino acids in length. Consistent with early embryonic lethality in knock-out mice, ATR-X patients carry hypomorphic alleles rather than complete loss of function mutations. There are no patients with large deletions in the gene, but several patients carry a nonsense mutation or frameshifting indel in early exons, which at first glance would be predicted to truncate large portions of the protein or deplete the transcript if targeted for nonsense mediated decay (NMD). Instead, these “early truncating” mutations can be partially rescued by low levels of translation to produce protein lacking the extreme N-terminus from a downstream ATG, as demonstrated for R37X (Howard et al., 2004). An alternative rescue mechanism is splicing out mutated exons, converting a truncating mutation into a short in-frame deletion, again with low production levels (Gibbons et al., 2008). Notably, the resulting ATRX alleles with these “early truncating” mutations do not impact known functional domains. In contrast, missense mutations (and in-frame indels within exons that cause the gain/loss of up to 22 amino acids), cluster in two functional domains: the N-terminal chromatin binding ADD (ATRX-DNMT3-DNMT3L) domain and the C-terminal ATPase domain (Gibbons and Higgs, 2000). Quantification of protein levels in EBV-transformed patient lymphocytes has helped determine the extent to which these mutations impact protein stability versus functionality. Protein levels in these mutants range from 7% to 51% of controls, and functional analysis of the more stable mutants has demonstrated reduced heterochromatin binding or chromatin remodelling activity (Argentaro et al., 2007; Iwase et al., 2011; Mitson et al., 2011).

A mouse line that carries a constitutive hypomorphic mutation should more effectively model ATR-X syndrome. Mice carrying a deletion of exon 2 model the “early truncating” mutations, expressing low levels of N-terminally truncated protein. These mice have been shown to recapitulate several aspects of the patient syndrome, including neurological defects and reduced brain size (Nogami et al., 2011; Shioda et al., 2011). Here, we produced the first model carrying a true ATR-X patient mutation. We chose the ADD mutant R246C (R245C in mice) as it is by far the most common patient mutation. Since R246C protein has previously been shown to be relatively stable but have impaired binding to heterochromatin, we predicted that this model would shed light on the importance of the ADD domain (Argentaro et al., 2007; Iwase et al., 2011). The mutant protein is sufficient to rescue the apoptotic cell loss phenotype that leads to death by 48 hours of age, described in mice lacking ATRX in the forebrain (Bérubé et al., 2005; Seah et al., 2008). The mice carrying R245C survive to adulthood, developing several neurological defects and morphological features reminiscent of ATR-X syndrome, such as craniofacial dysmorphism and microcephaly. This represents the patient condition, where microcephaly is acquired after birth (Gibbons, 2006). Intriguingly, the mutant protein is relatively stable (∼60% of controls) during embryonic development but declines dramatically to ∼10% of that seen in controls by 9 weeks of age, suggesting that instability may contribute more to the pathogenicity of R246C and other “stable” mutations than previously thought. We anticipate that the new R245C mouse model will be adopted by the field to further our understanding of the molecular mechanisms underlying ATR-X syndrome and test future therapeutic avenues.

## Results

### Production of *Atrx^R245C/y^* knock-in mice

There are >250 patients with ATR-X syndrome and >100 causative mutations have been identified (Gibbons et al., 2008). Strikingly, the missense mutations and short in-frame indels fall almost entirely into two clusters: the chromatin-binding ADD (ATRX-DNMT3-DNMT3L) domain and the ATPase domain (Figure 1A). The ADD and ATPase clusters represent 43% and 32% of cases, respectively and the most common patient mutation, R246C, is found in 18% of cases (Figure 1B). We therefore selected R246C as the most suitable candidate for a patient mutation mouse model of ATR-X syndrome. Knock-in mice were produced by electroporation of C57BL/6J zygotes with RNP (Cas9 protein and synthetic sgRNA) and a 142 nt donor template containing the desired mutation (c.733C>T; p.R245C, mouse homolog of R246C) and a silent PAM-abolishing mutation (c.729C>T; p.I243I) (Figure 1C). We generated eight hemizygous male founders and one homozygous female founder carrying both mutations, and successfully bred five of the males. We established lines from two founders and carried out quality control assays on N1 heterozygous females and N2-3 males. Both lines carried the desired mutations, had no additional insertion of the donor template, and pups were born at the expected Mendelian ratio (Figure S1). We chose line 1 for all future experiments. To best model ATR-X patients, hemizygous males were used for all analyses.

**Figure 1.**
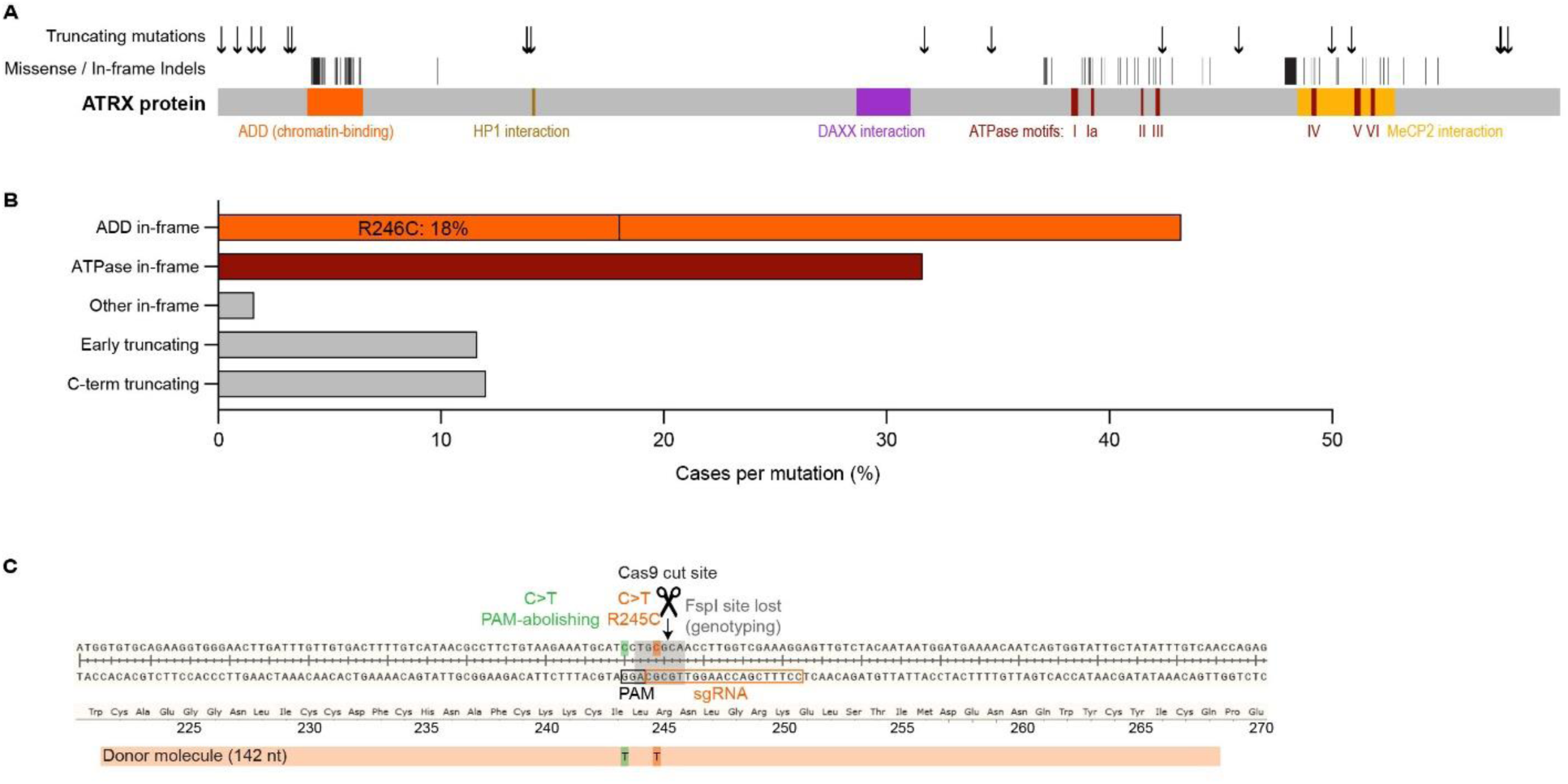
Production of *Atrx^R245C/y^* knock-in mice. **A.** Schematic showing ATRX patient mutations, divided into truncating and in-frame mutations (missense and short indels). **B.** Frequency of ATR-X syndrome-causing mutations identified in patients, separated by category (ADD in-frame, ATPase in-frame, other in-frame, “early truncating”, and C-terminal truncating). **C.** *Atrx^R245C/y^* mice were produced by Cas9-mediated editing in C57BL/6J embryos. Schematic of exon 9 of the mouse *Atrx* gene. A double-strand break was introduced by sgRNA-guided Cas9 and a 142 nt ss oligo containing the desired mutation (c.733C>T; p. R245C) and a PAM-abolishing silent mutation (c.729C>T p. I243I) was incorporated. Mice were routinely genotyped using FspI digestion, which cleaves the WT PCR product.

### The R245C mutation impacts ATRX protein stability and binding to heterochromatin

Western blot analysis of ATRX protein levels in cortical tissue at birth (P0.5), found that the mutant protein was decreased to ∼60% of wild-type controls (Figure 2A). This is consistent with a previous report that ATRX[R246C] is relatively stable (38% of controls) in patient lymphocytes (Argentaro et al., 2007). Surprisingly, repeating this analysis in mouse cortical tissue at 9 weeks of age found that the mutant protein was dramatically depleted to ∼10% of wild-type levels (Figure 2B). This is unlikely to be a result of reduced *Atrx* transcription, as mRNA levels were only minorly decreased (∼80% of controls) at both timepoints (Figure 2C). We therefore investigated protein stability with the translation-inhibiting drug, cycloheximide (CHX), in cultured cortical neurons derived from *Atrx^R245C/y^* and wild-type embryos. After three or 8 hours of CHX treatment, the wild-type protein was present at similar levels as in DMSO-treated controls, but the mutant protein was depleted to 68% or 60% (Figure 2D), indicating that the R245C mutation reduces stability. Analysis of adult spleen tissue found that ATRX[R245C] levels are similarly decreased to ∼15% of wild-type levels (Figure 2E), demonstrating that the postnatal decrease in mutant ATRX levels was not unique to the brain. Functionality of the heterochromatin binding domain can be assessed in mouse cells and tissues by localisation of ATRX at H3K9me3-rich pericentromeric foci (visualised as DAPI bright spots). Overexpression of human ATRX in cultured mouse fibroblasts revealed punctate staining for wild-type protein but diffuse staining for R246C protein (Iwase et al., 2011). Here, we found that the R245C mutation impacts recruitment of endogenous ATRX to heterochromatic foci, detected using immunofluorescence staining in mouse brain tissue at P0.5 (Figure 2F). We conclude that impaired functionality is likely to be the main contributing factor to R245C pathogenicity during embryonic development, when ATRX[R245C] is relatively stable, but very low mutant protein levels resulting from reduced stability may be the dominating factor contributing to the pathogenicity of this mutation in adulthood.

**Figure 2.**
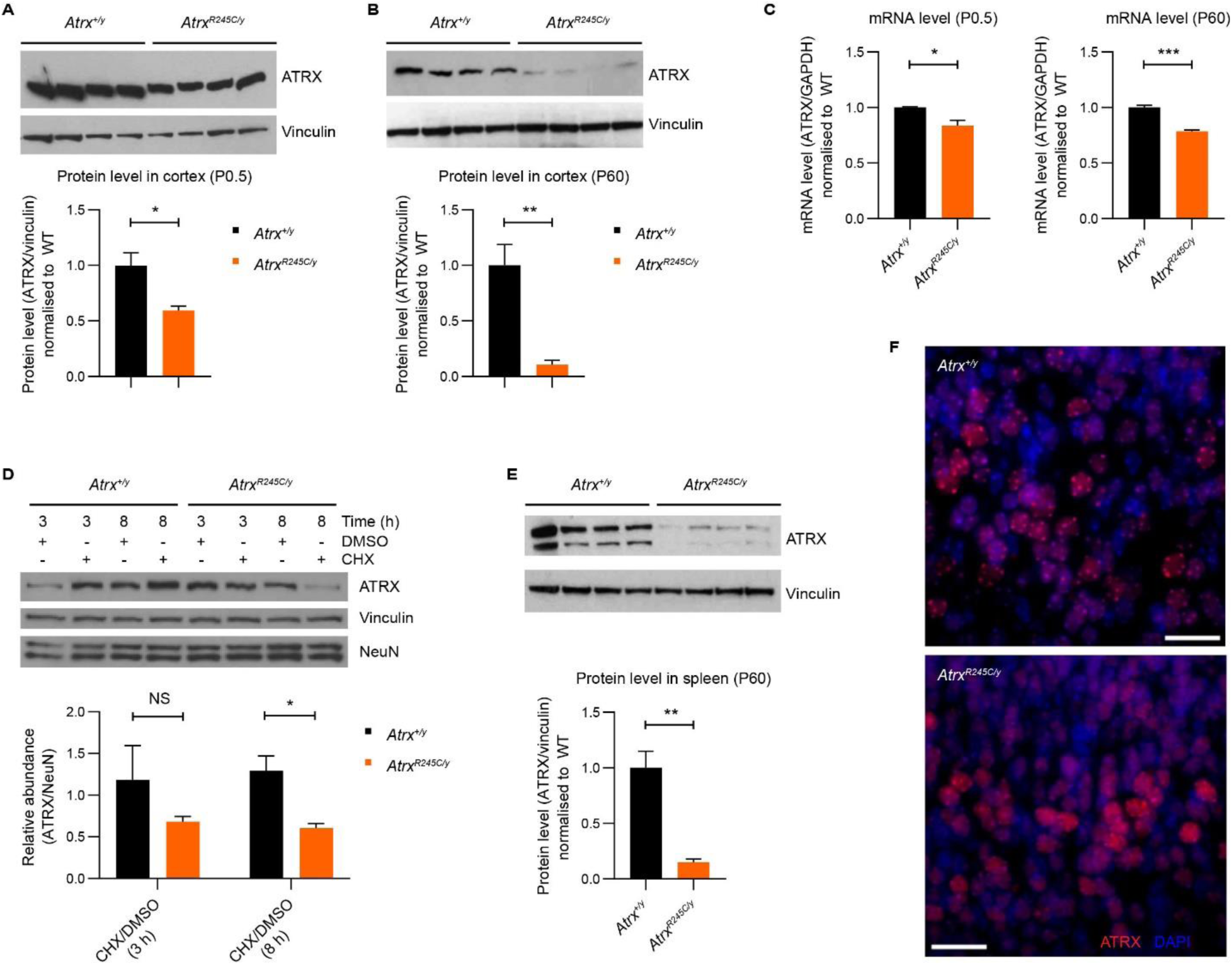
The R245C mutation impacts ATRX protein stability and binding to heterochromatin. **A-B.** Western blot analysis of ATRX[R245C] protein levels in cortex of *Atrx^R245C/y^* males at (**A**) P0.5 and (**B**) P60, compared to wild-type controls (n = 4 per genotype for each). Quantification (below): R245C protein is present at 59.5% and 10.7% of normal levels at P0.5 and P60, respectively. Graphs show mean ± S.E.M. and genotypes were compared using t-tests: P0.5 * P = 0.014; P60 ** P = 0.004. **C.** qPCR analysis of *Atrx*[R245C] transcript levels in cortex at P0.5 (left) and P60 (right), compared to wild-type controls (n = 3 per genotype for each). mRNA is expressed at 83.8% and 78.4% of WT levels at P0.5 and P60, respectively. Genotypes were compared using t-tests: P0.5 * P = 0.03; P60 *** P = 0.0009. **D.** Cortical neurons at DIV7 derived from *Atrx^+/y^* and *Atrx^R245C/y^* E17.5 embryos were treated with DMSO vehicle or 100 μM Cycloheximide (CHX) for 3 or 8 hours. Quantification (below) of the relative levels of ATRX (normalised to NeuN) in cells treated with CHX vs DMSO (3 replicates). Graph shows mean ± S.E.M. and genotypes were compared using t-tests: 3 h P > 0.5; 8 h * P = 0.02. **E.** Western blot analysis of ATRX[R245C] protein levels in spleen at P60, compared to wild-type controls (n = 4 per genotype). Quantification (below): R245C is present at 15.2% of normal levels. Graph shows mean ± S.E.M. and genotypes were compared using a t-test: ** P = 0.0014. **F.** Immunofluorescence staining of ATRX (red) in the cortex at P0.5. Nuclei are stained with DAPI (blue). Scale bar: 20 µm.

### *Atrx^R245C/y^* mice have reduced body size, brain weight and a craniofacial phenotype

The morphological features of ATR-X syndrome can be evident in children from a very early age and include short stature, postnatal microcephaly, facial dysmorphism, hypotonia, and genital abnormalities (Gibbons, 2012). As such, we performed a variety of morphological analyses on the *Atrx^R245C/y^* mice to assess similarity to the human phenotype. *Atrx^R245C/y^* mice were lighter than wild-type littermate controls from weaning until at least one year of age (Figure 3A). This was not due to a difference in lean/fat ratio, measured by DEXA at P21 (Figure S2A). X-ray analysis showed that mutant mice had smaller skeletons, with reduced body length in early (14 weeks) and late (52 weeks) adulthood (Figure 3B). This is consistent with short stature in ATR-X syndrome (Gibbons, 2012). As patients have acquired microcephaly (Gibbons, 2006), we measured brain weight at 9 weeks, which was decreased in the mutants (Figure 3C). Facial dysmorphism is highly characteristic of ATR-X patients (Gibbons, 2012), so we assessed facial appearance in the mice (blind to genotype), observing that 11/19 (58%) of mutants in the initial cohort had visibly shortened snouts (Figure 3D), compared to 2/20 (10%) of wild-type littermate controls. Quantification using X-ray imaging revealed a significant decrease in cranial length at 14 weeks (Figure 3E-F). However, when skull shape was analysed by determining the cranial index (cranial width/cranial length x 100), this difference was no longer significant (Figure 3G). Mutant animals had a shorter zygomatic bone length (Figure 3H) and analysis of shape using “zygomatic index” (zygomatic width/zygomatic length x 100), confirmed this difference (Figure 3I). These findings were reproduced in an independent cohort, aged 40 weeks (Figure 3F-I).

**Figure 3.**
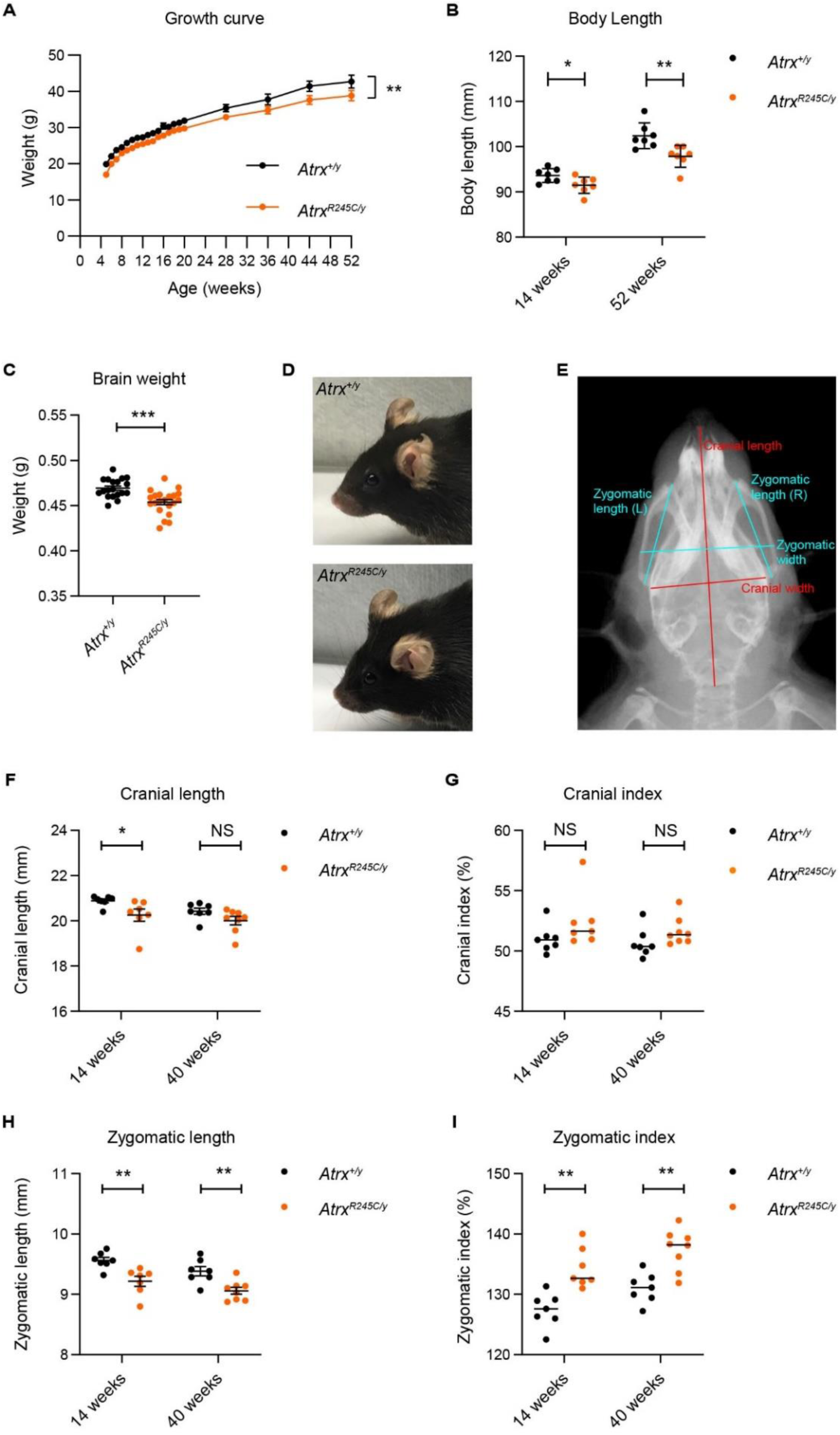
*Atrx^R245C/y^* mice have reduced body weight and craniofacial defects *Atrx^R245C/y^* mice have reduced body weight and craniofacial defects. **A.** Growth curve showing body weight from 5-52 weeks of age (WT n = 13; *R245C* n = 13). Graph shows mean ± S.E.M. and genotypes were compared using repeated measures ANOVA: ** P = 0.002. **B.** Body length (nose to base of tail) measured from X-ray analysis (WT n = 7; *R245C* n = 7). Genotypes were compared using t-tests: 14 weeks * P = 0.03; 52 weeks ** P = 0.007. **C.** Brain weight at 9 weeks (WT n = 19; *R245C* n = 21). Genotypes were compared using a t-test: *** P = 0.0002. **D.** Representative photographs of the shortened snout phenotype observed in ∼60% of *Atrx^R245C/y^* mice (lower), compared to WT controls (upper). **E.** Example ventral X-ray of a WT mouse showing how cranial length, cranial width, zygomatic length (mean of left and right) and zygomatic width were measured. Cranial index [also known as cephalic index] = cranial width/cranial length x 100 and zygomatic index = zygomatic width/zygomatic length x 100. **F-I.** X-ray analysis quantifying cranial length (**F**) cranial index (**G**), zygomatic length (**H**) and zygomatic index (**I**) at 14 weeks (WT n = 7; *R245C* n = 7) and 40 weeks (WT n = 7; *R245C* n = 8). Graphs of cranial length and zygomatic length show mean ±S.E.M. and genotypes were compared using t-tests: cranial length at 14 weeks * P = 0.47; at 40 weeks P = 0.099; zygomatic length at 14 weeks ** P = 0.004; at 40 weeks ** P = 0.004. Graphs of cranial index and zygomatic index show median. Cranial index: genotypes were compared using KS tests: 14 weeks P = 0.21; 40 weeks * P = 0.03. Zygomatic index: genotypes were compared using t-tests: 14 weeks ** P = 0.001; 40 weeks ** P = 0.001.

ATR-X patients have neonatal hypotonia (Gibbons, 2012) and mice lacking ATRX in skeletal muscle have reduced muscle mass and function at weaning, which is normalised by adulthood (Huh et al., 2012). In contrast, tibialis anterior (TA) muscle weight in *Atrx^R245C/y^* mice at P22 was proportional to body weight (Figure S2B). Similarly, grip strength, used as a measure of muscle strength, was also proportional to body weight in these animals (Figure S2C). We assessed motor function more extensively in early adults (9-14 weeks of age) and found no differences in grip strength (Figure S2C), minor differences in gait (Figure S2D), but normal performance on the accelerating rotarod (Figure S2E). Since mice lacking ATRX in skeletal muscle show reduced performance with chronic exercise (Huh et al., 2017), we exercised an independent cohort of animals on a treadmill with an increasingly steep decline for four training days, followed by three trial days. Very few mice were unable to complete trials and this was comparable between genotypes (Figure S2F). After chronic exercise, we saw no evidence of TA muscle damage from permeation of Evan’s blue dye and no fibrosis in either genotype (Figure S2G). Additionally, we observed no difference in proportion of smaller muscle fibres in the mutants (Figure S2H), a sign of muscle regeneration. Analysis of older adult mice also found no age-related decline in grip strength (Figure S2C) and decreased stride length (Figure S2I) is likely linked to smaller body size. Although 3/13 aged mutant mice failed to run on the treadmill used for gait analysis, an independent cohort displayed normal performance on the accelerating rotarod (Figure S2J). Kyphosis is thought to be connected with muscle defects in patients since it was observed in muscle cKO mice (Huh et al., 2012). However, the kyphotic index was unaltered in *Atrx^R245C/y^* mice at weaning or in early/late adulthood (Figure S2K). Overall, we were unable to detect muscle defects in *Atrx^R245C/y^* mice.

We did not observe any morphological differences in genitalia in our *Atrx^R245C/y^* mice and at least five hemizygous male founders and all tested males from both line 1 (n=3) and line 2 (n=3) were fertile. While genital abnormalities are frequently reported in ATR-X syndrome patients (70-80%), they are most severe in those with C-terminal truncations (Gibbons et al., 2008; Stevenson, 2020). Reduced expression of alpha-globin (*HBA*), resulting in alpha-thalassemia, is detected in ∼75% of ATR-X patients (Stevenson, 2020). Reduction in *HBA* expression, and therefore severity of thalassemia, correlates with the length of a G-rich VNTR repeat within the nearby HBAZ(ps) pseudogene (Law et al., 2010). Analysis of the mouse *Hba* locus revealed that this repeat is not conserved across species (Figure S3A). Accordingly, *Hba* expression and haematology are unaffected in *Atrx^R245C/y^* mice (Figure S3B-G). Altogether, the *Atrx^R245C/y^* mice recapitulate several morphological features of ATR-X syndrome, including short stature, postnatal microcephaly and facial dysmorphism, but this model lacks hypotonia, genital abnormalities and alpha-thalassemia.

### *Atrx^R245C/y^* mice display neurobehavioural phenotypes

As patients with ATR-X syndrome have mild to profound intellectual disability, we performed neurological tests on the *Atrx^R245C/y^* mice. Behavioural assays are typically done using young adults (9-14 weeks), when ATRX[R245C] protein levels were decreased to ∼10% in the brain (Figure 2B). All reflexes were normal based on SHIRPA analysis, but *Atrx^R245C/y^* mice had lower spontaneous activity than wild-type littermate controls (Figure 4A). The Open Field test, which measures activity for a longer period of time, however, found normal exploratory behaviour (Figure S4A). Mutants also displayed normal levels of anxiety with regards to open space (Figure S4B), light (Figure S4C) and height (Figure S4D). These findings allowed us to assess more complex behaviours, without the compounding effects of altered activity or anxiety. We saw no genotypic differences in motor learning over three days of testing on the accelerating rotarod, with both genotypes displaying daily improvement (Figure S4E). Spatial memory was assessed over four days in the Barnes Maze, where wild-type mice locate the hidden escape hole with increasing ease, but mutants displayed impaired performance, which was significantly different from controls on three of four days (Figure 4B). Social Interaction was assessed in the three chamber test, where wild-type controls displayed a significant preference for the novel stranger mouse, but this was lost in the mutants (Figure 4C). Given the results from the Barnes Maze and Social Interaction test, we asked whether defects in spatial memory and novelty preference affect the performance of *Atrx^R245C/y^* mice in the Y-maze. However, over the 8-minute duration of this test, we saw no difference in Alternation Index between genotypes (Figure S4F). Lastly, in the Fear Conditioning paradigm, *Atrx^R245C/y^* mice showed impaired contextual but unaltered cued fear memory (Figure 4D; S4G). We repeated some of these tests in late adult mice (around one year of age) and found a slight increase in hypoactivity (Figure S5A-B) but no late-onset memory defects (Figure S5C-F). Overall, our findings that *Atrx^R245C/y^* mice display neurological phenotypes was do not worsen with age are consistent with the classification of ATR-X syndrome as a developmental rather than degenerative condition.

**Figure 4.**
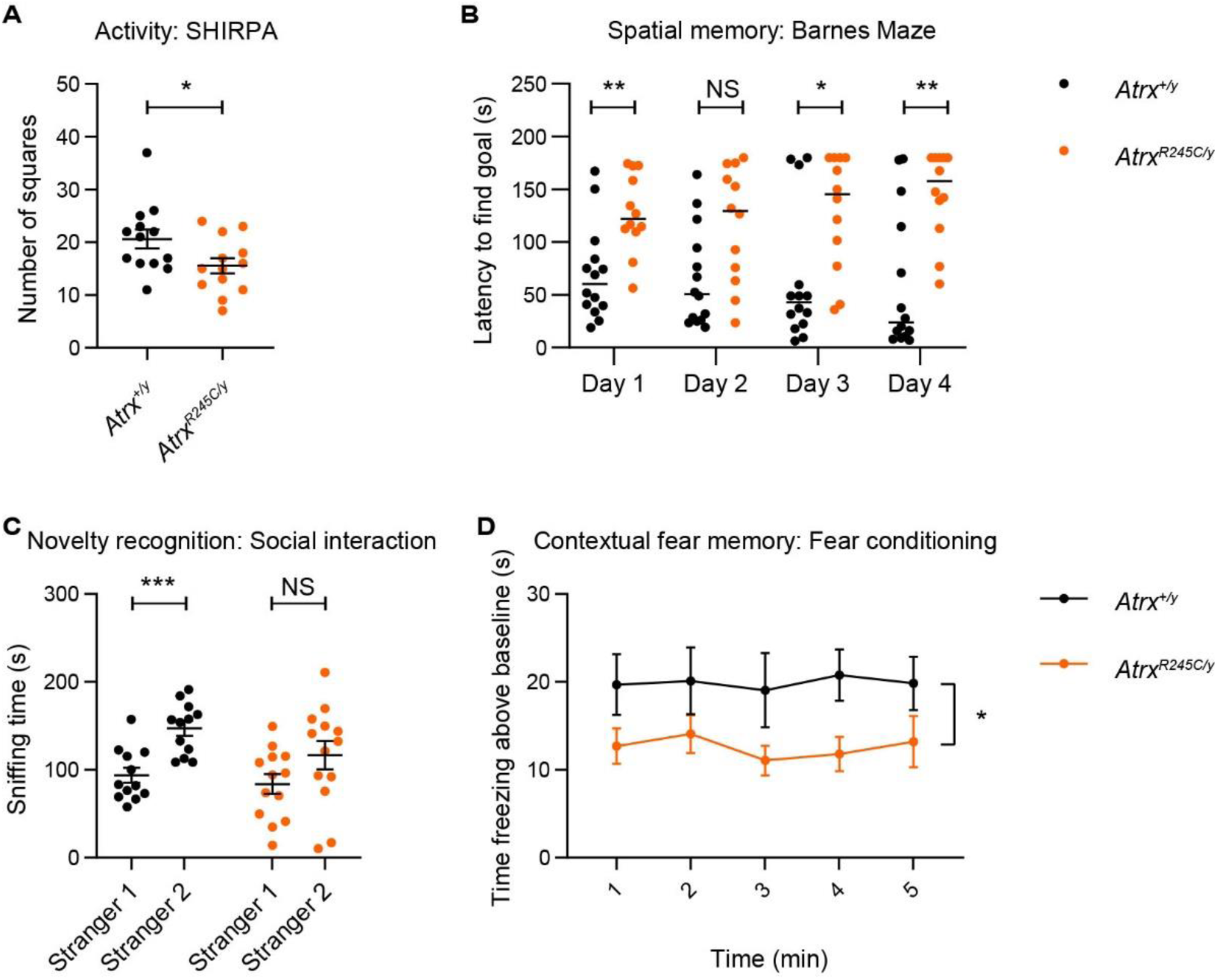
*Atrx^R245C/y^* mice have neurological defects. **A.** Spontaneous activity assessed by number of squares entered in 30 s at 9 weeks (WT n = 13; *R245C* n = 13). Graphs show mean ± S.E.M. and genotypes were compared using a t-test: * P = 0.04. **B.** Mean latency to find the escape hole in the Barnes maze test in four trials was calculated for each of the 4 days of the experiment at 12 weeks (WT n = 14; *R245C* n = 12). Graphs show median and genotypes were compared using KS tests: day 1 ** P = 0.004, day 2 P > 0.05, day 3 * P = 0.014, day 4 ** P = 0.0096. **B.** Time spent sniffing a cylinder container a familiar (Stranger 1) or novel (Stranger 2) mouse in the three-chamber social interaction test at 10 weeks (WT n = 12; *R245C* n = 13). Graph shows mean ± S.E.M. and Stranger 1 vs 2 sniffing times were compared using paired t-tests: WT *** 0.0004, *R245C* P > 0.05. Genotypes were compared using 2-way ANOVA: P = 0.09. **D.** Contextual fear conditioning analysis at 13 weeks (WT n = 14; *R245C* n = 13). Time spent freezing (minus baseline determined before shock) when returned to the same contextual environment 24 h after receiving a foot shock. Graph shows mean ± S.E.M. and genotypes were compared using repeated measures ANOVA: * P = 0.025.

### Cerebellar volume is reduced in *Atrx^R245C/y^* mice

Perinatal lethality in mice lacking ATRX in the forebrain is associated with apoptotic cell death, resulting in reduced brain size, reduced cortical density and the replacement of the dentate gyrus of the hippocampus with a small mass of disorganised cells (Bérubé et al., 2005; Huh et al., 2016; Seah et al., 2008). In contrast, brains of the *Atrx^R245C/y^* mice show comparable morphology to controls at birth (Figure 5A) and staining of cortical layer markers demonstrates that cellular differentiation in the cortex is unaffected and cell numbers are equivalent (Figure 5B-D). These findings are consistent with normal head circumference at birth in ATR-X syndrome patients (Gibbons, 2006).

**Figure 5.**
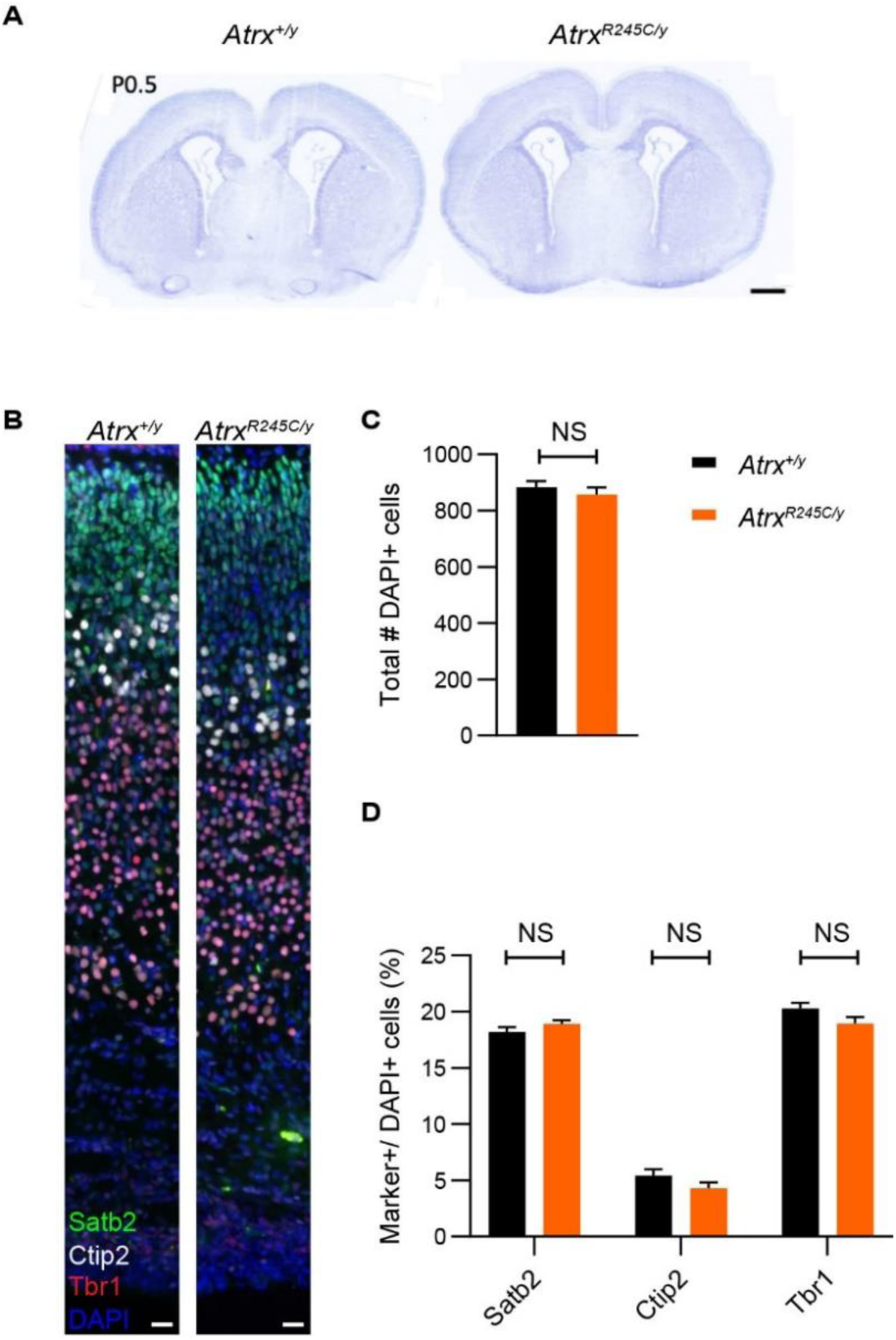
Embryonic cortical development is unaffected in *Atrx^R245C/y^* mice. **A.** Representative images of Nissl-stained coronal section from WT and R245C mice at P0.5. Scale bar: 500 µm. **B.** Representative images of P0.5 brain sections stained for cortical layer markers Tbr1 (red, layer VI); Ctip2 (white, layer V), and Satb2 (green, layers II-V). Nuclei are counterstained with DAPI (blue). Scale bar: 20 µm; n=4. **C.** Total number of DAPI-positive cells in the cortex (n= 6 biological replicates per genotype). **D.** Quantification of layer marker+ cells within the neocortex (n= 6 biological replicates per genotype). Graphs show mean ± S.E.M. and genotypes were compared using t-tests: all NS P > 0.05.

We therefore focused on the early adult timepoint, when *Atrx^R245C/y^* mice displayed neurological phenotypes (Figure 4). MRI analysis revealed a slight downward trend in total brain volume at 9 weeks (Figure 6A; -1.2% change, P = 0.14). Notably, this difference was not as robust as the decrease in brain weight, measured at the same age in a separate cohort (Figure 3C; -3.3% change, *** P = 0.0002).

**Figure 6.**
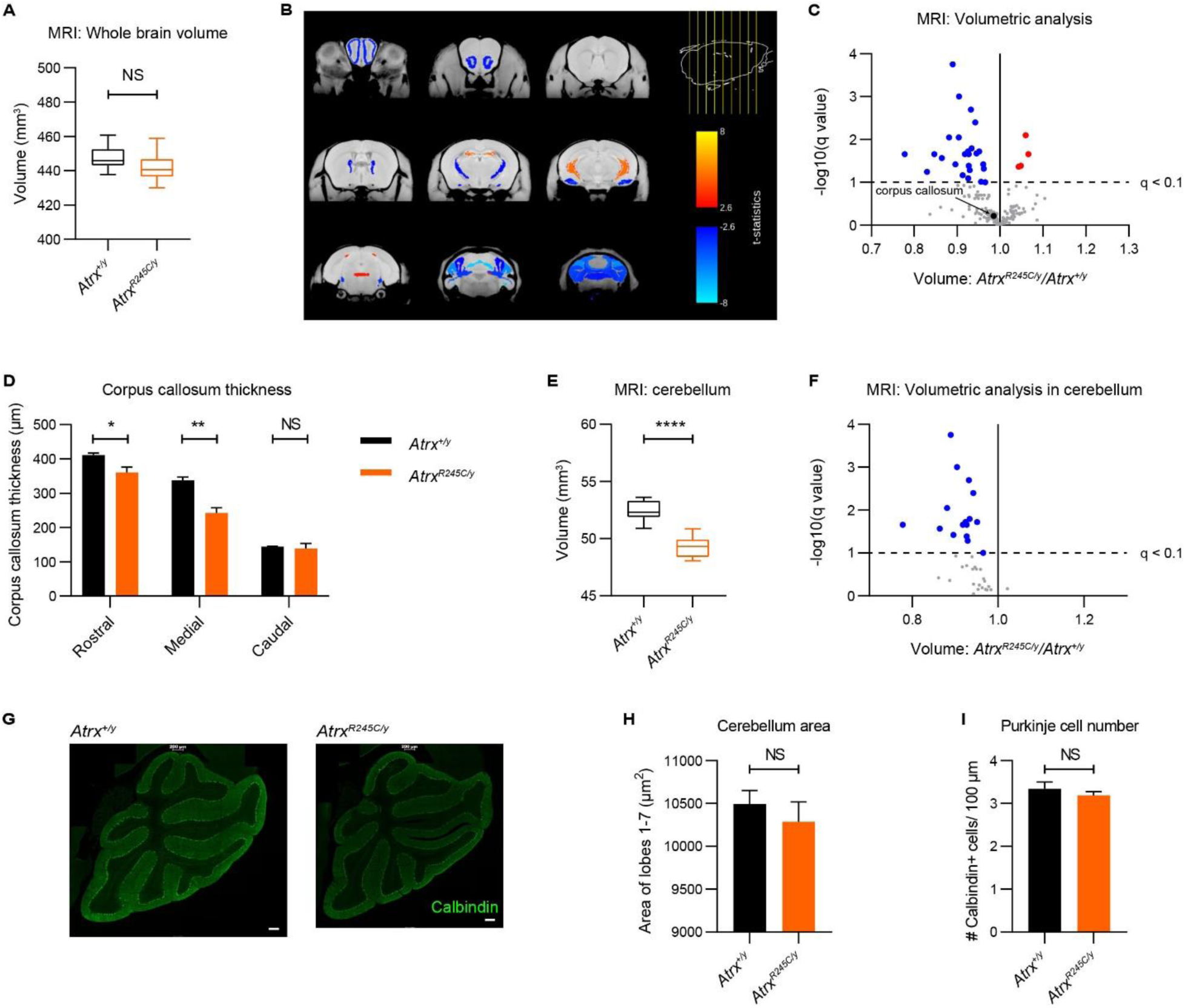
Cerebellar volume is reduced in *Atrx^R245C/y^* mice. **A.** Whole brain volume analysed by MRI at 9 weeks (WT n = 10; *R245C* n = 12). Graph shows interquartile ranges and genotypes were compared using a t-test: P > 0.05. **B.** Coronal sections showing significant changes in absolute volume in *Atrx^R245C/y^* mouse brains (coloured regions indicate significant changes at q < 0.1). **C.** Volcano plot of all 183 brain regions. Significantly changed (q < 0.1) regions are highlighted in red (increased) and blue (decreased). The corpus callosum is highlighted in black. **D.** Quantification of corpus callosum thickness measured from rostral, medial and caudal coronal brain sections stained with myelin-associated glycoprotein (MAG) at P40 (WT n = 4; *R245C* n = 4). Graph shows mean ± S.E.M. and genotypes were compared using t-tests: rostral * P = 0.023; medial ** P = 0.002; caudal P > 0.05. **E.** Total cerebellar volume analysed by MRI at 9 weeks (WT n = 10; *R245C* n = 12). Graph shows interquartile ranges and genotypes were compared using a t-test: **** P < 0.0001. F. Volcano plot (as in Figure 6C) showing volumetric analysis of the 39 cerebellar regions. **G.** Representative images of cerebellar sections from WT and R245C mice at P40, stained with Calbindin, a marker of Purkinje neurons (green). Scale bar: 200 µm. **H-I.** Quantification the total area of the cerebellar section (**H**) and the number of Calbindin+ cells per 100 µm across lobes I-VII (**I**) (WT n = 3; *R245C* n = 3). Graphs show mean ± S.E.M. and genotypes were compared using t-tests: both NS P > 0.05.

Reduced cortical thickness was not a contributing factor to microcephaly in adult mice, as this was only decreased in the ventral orbital cortex and was slightly, but significantly, increased in 8 out of the 38 cortical regions analysed (Figure S6A). Published MRI analysis of patients’ brains found a variety of grey and white matter abnormalities, most frequently non-specific brain atrophy (Wada et al., 2013). This was thought to be a result of reduced neuronal or glial production in the postnatal period rather than due to degeneration, though it has been reported to be progressive in two cases (Lee et al., 2015; Wada et al., 2013). While affected brain regions vary among patients, there are several reports of partial or complete agenesis of the corpus callosum (Gibbons, 2006; Thienpont et al., 2007; Wada et al., 2013). To determine whether any regions are disproportionately smaller in the mutant mice, we performed comparative volumetric analysis on 183 brain subregions. Using a significance threshold of q < 0.1 (after correction for multiple comparisons by the false discovery rate), 26 regions were smaller, and four regions were larger in mutants than in wild-type littermate controls (Figure 6B-C), consistent with reduced brain size. This analysis did not detect a significant decrease in corpus callosum volume (Figure 6C; S6B). However, quantification of corpus callosum thickness in rostral, medial and caudal histological sections revealed that it was significantly thinner in mutant mice in rostral (−12.3% change, * P = 0.02) and medial (−28.0% change, ** P = 0.002) regions (Figure 6D; S6C). Intriguingly, the MRI data revealed volumetric differences in regions of the cerebellum (Figure 6B). Total cerebellar volume was significantly decreased (−6.0% change, **** P < 0.0001; Figure 6D) and 17/39 regions in the cerebellum were significantly smaller (Figure 6E; Figure S6D-E). We therefore asked whether reduced cerebellar volume was due to reduced cell number. Given that the cerebellum undergoes a period of rapid expansion in early postnatal development (Leto et al., 2016), it could be particularly susceptible to increased replicative stress in the mutants. By analysis of cerebellar size using immunofluorescence staining, we observed a slight but not significant decrease in the area of cerebellar sections (Figure 6G-H), reflecting the volumetric analysis. This was not accompanied by a significant decrease in the number of Purkinje (calbindin+) neurons (Figure 6I), suggesting that reduced cerebellar volume does not result from reduced proliferation or increased cell death.

Mice lacking ATRX in postnatal forebrain excitatory neurons (via *CaMKII-Cre* mediated deletion) display similar neurobehavioural defects to those we observed in *Atrx^R245C/y^* mice, including impaired spatial memory and contextual fear memory (Tamming et al., 2020). *CaMKII-cKO* mice have reduced whole brain volume and differences in hippocampal neuroanatomy, consistent with the role of the hippocampus in memory and learning. MRI analysis showed that the total hippocampal volume is unaffected in *Atrx^R245C/y^* mice (Figure S7A). We next compared our MRI volumetric analysis for hippocampal regions (*Atrx^R245C/y^* vs wild-type controls) with those reported for the *CaMKII-cKO* mice (vs wild-type controls). Of the seven regions affected in the *CaMKII-cKO* mice, three were also affected in the *Atrx^R245C/y^* mice, but in the opposite direction (Figure S7B-D). We therefore found no correlation between the two models. We further analysed hippocampal structure by immunofluorescence staining and observed no differences in morphology (Figure S7E). There were no differences in the proportion of total mature neurons (NeuN+) in the CA1, CA3 and dentate gyrus. Additionally, there were no differences in the total number of cells or the proportion of neuronal progenitor cells (doublecortin, DCX+) and interneurons (Calretinin+) in the dentate gyrus (Figure S7F-H). In summary, changes in brain structure observed in *Atrx^R245C/y^* mice are distinct from those previously reported in *Foxg1-* and *CaMKII-cKO* models. Our brain morphology observations show overlap with the postnatal microcephaly and corpus callosum hypoplasia described in patients. Interestingly, this is the first report of reduced cerebellar volume in *Atrx*-mutant mice.

### *Atrx^R245C/y^* neurons have reduced dendritic branching

To further address the cause of reduced brain size, we performed Sholl analysis on cultured hippocampal neurons derived from *Atrx^R245C/y^* mice and wild-type controls (Figure 7A-B). Axon length was unchanged between genotypes, but total dendritic length was decreased in the mutant cells (Figure 7C-D). This is consistent with a decrease in dendritic branching (Figure 7E). It is therefore possible that microcephaly in *Atrx^R245C/y^* mice and ATR-X syndrome patients in part results from smaller, less complex cells.

**Figure 7.**
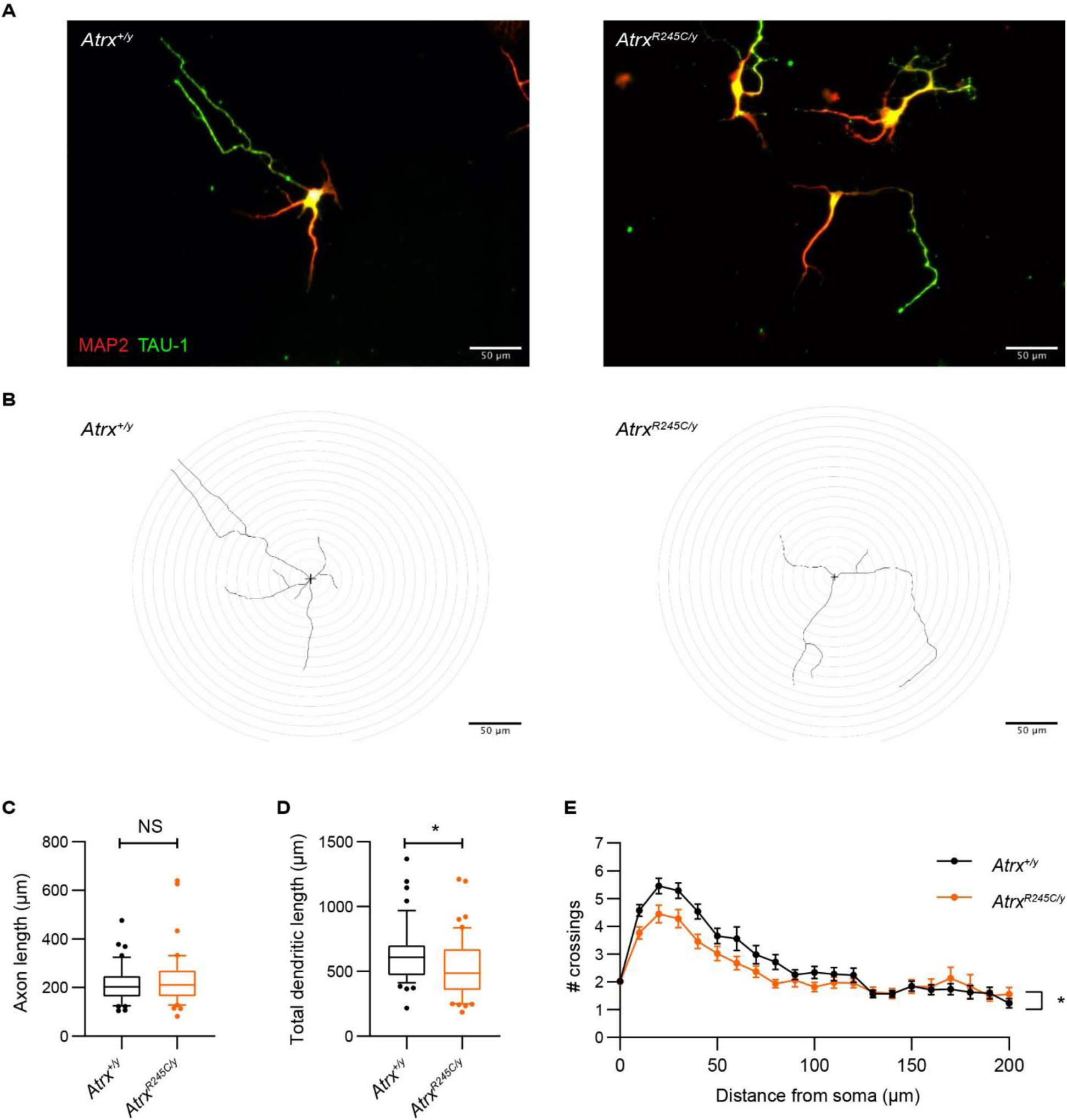
*Atrx^R245C/y^* neurons have reduced dendritic length and branching. **A.** Hippocampal neurons derived from *Atrx^+/y^* and *Atrx^R245C/y^* mice at E17.5 (DIV5) stained with MAP2 (red) and TAU-1 (green). **B.** Representative tracings of the neurons shown in (**A**), used for Sholl analysis. **C-D.** Axon length (**C**) and total dendritic length (**D**) of the neurons (WT n = 49; *R245C* n = 50). Graphs show interquartile ranges (whiskers 10-90 percentile) and genotypes were compared using KS tests: axon length P > 0.05; total dendritic length * P = 0.043). **E.** Sholl analysis of dendritic branching in the same cells. Graph shows mean ± S.E.M. for each distance, and genotypes were compared using Mixed-Effects analysis: * P = 0.015.

## Discussion

We present a mouse mutant containing the first engineered patient mutation as a novel model of ATR-X syndrome, carrying the most frequently occurring pathogenic variant: R246C. These mice recapitulate several morphological and neurological aspects of the disorder, including reduced body size, reduced brain weight, craniofacial defects and impaired brain function. These are reminiscent of the short stature, microcephaly, facial dysmorphism and intellectual disability described in patients (Gibbons, 2012; Stevenson, 2020). Overall, we found strong phenotypic overlap between the *Atrx^R245C/y^* mice and the “early truncating” model lacking exon 2, which were also characterised on C57BL/6J (Nogami et al., 2011; Shioda et al., 2011). Specifically, both models have reduced body weight, brain weight, impaired spatial memory in the Barnes Maze, impaired novelty preference (analysed using the Social Interaction test for *Atrx^R245C/y^* and the Novel Object Recognition test for *Atrx^Δex2/y^*) and impaired contextual fear memory. While defects in spatial memory and/or novelty preference displayed by the *Atrx^Δex2/y^* mice were also evident as decreased Alternation index in the Y-maze, this result wasn’t reproduced in the *Atrx^R245C/y^* mice. Historically, patients with different mutations in the same gene have often been classed as having distinct conditions, sometimes with unique names. This has been the case for ATR-X syndrome: for example, some patients with the “early-truncating” mutation R37X were reported to have Chudley-Lowry syndrome (Abidi et al., 2005). The presence of a phenotypic signature shared by both mouse models carrying hypomorphic alleles of *Atrx* supports the move in the rare disease field towards categorising genetic diseases by the affected gene.

This is the first report of a craniofacial phenotype (shortened snout) in a mouse model with reduced ATRX function. Intriguingly, transgenic mice overexpressing ATRX have a similar phenotype (Bérubé et al., 2002). Further work is needed to determine the mechanism by which ATRX dosage affects skull morphology and whether the penetrance of the shortened skull phenotype could be modified by backcrossing to other strains.

We found that the *Atrx^R245C/y^* mice did not recapitulate the alpha-thalassemia, genital abnormalities, or muscle hypotonia described in patients. The absence of the first two features can be explained by the lack of the G-rich Ψζ VNTR repeat, previously linked to ATRX-mediated alpha-globin gene regulation, at the mouse locus (Law et al., 2010) and the connection between genital abnormalities and C-terminal truncating mutations (Gibbons et al., 2008; Stevenson, 2020). The absence of muscle defects in *Atrx^R245C/y^* mice was surprising, as deletion of *Atrx* in skeletal muscle (via *Myf5-Cre* mediated excision) resulted in neonatal hypotonia and impaired recovery after chronic exercise-induced damage (Huh et al., 2012, 2017). It is possible that the hypomorphic allele (R245C) retains enough functionality to rescue these phenotypes on C57BL/6J, but they could be detectable on another background strain. In support of this, the commonly used model of Duchenne Muscular Dystrophy (DMD), *Dmd^mdx^*, which carries a truncated *DMD* allele, is minimally affected on several background strains including C57BL/10 but displays severe muscular dystrophy on DBA/2J (Fukada et al., 2010). The ameliorating effect of the C57BL/6 background was overcome by removing residual DMD activity, as demonstrated in a recently published *DMD* knockout model, *Dmd^em1Rcn^*(Wong et al., 2020). This could also explain why the muscle phenotypes displayed by the *Atrx Myf5-cKO* mice were not detected in *Atrx^R245C/y^* mutants.

*Atrx^R245C/y^* mice can now be adopted by the field to further our understanding of the molecular mechanisms underlying ATR-X syndrome and to test potential therapeutic avenues. ATRX is believed to be a pleiotropic protein, due to its proposed roles in multiple molecular processes. Notably, these studies have been restricted to accessible patient cells (erythroblasts and lymphoblastoid cells), knock-out cell lines, and tissues derived from conditional knock-out mice (Bérubé et al., 2005; Garrick et al., 2006; Law et al., 2010; Truch et al., 2022). Our mice carry a constitutive hypomorphic mutation, representing the genetic status of all ATR-X patients. We have shown here that this model better represents patient disease pathology than *Atrx* conditional knock outs: *Atrx^R245C/y^* mice survive to adulthood, do not exhibit extensive neuronal cell loss and display neurobehavioural phenotypes. The reduction in neurite length and branching suggests that acquired microcephaly in patients could be related to abnormal neuronal morphology. Notably, the reduced brain weight phenotype in *Atrx^Δex2/y^* mice is accompanied by altered dendritic spine density (Shioda et al., 2011). Further work is needed to determine whether these models share these cellular phenotypes.

We found that ATRX[R245C] mutant protein is sufficient to rescue the neuronal death phenotype present in mice lacking ATRX in the forebrain. This is supported by a recent study using CRISPR-edited mouse ESC-derived neural progenitor cells (NPCs) where a cell death phenotype was observed in cells lacking ATRX, but not in those expressing a stable mutant ATRX with two amino acid changes in the ADD domain that specifically abolish heterochromatin binding (Bieluszewska et al., 2022). This raises the question of whether mutating the ADD domain can cause elevated replicative stress, as has been described in *Atrx-null* NPCs (Watson et al., 2013) and myoblasts (Huh et al., 2012). Having a well characterised patient-relevant mouse model of ATR-X syndrome means that we can investigate the role of ATRX protein in replication and other molecular processes in the most relevant tissues and developmental time points. This will help us understand which aspects of ATRX protein function are “missing” to result in ATR-X syndrome; and how residual functionality of a hypomorphic mutant rescues the more severe *Atrx*-null phenotypes. We predict that such studies will uncover molecular phenotypes shared across ATR-X syndrome-causing mutants, and those specific to ATRX[R245C] or ADD mutants. Dissecting the functionality of the ADD domain in heterochromatin maintenance, gene regulation and DNA damage repair is of particular interest as 43% of patients have mutations that lie within it. ATRX has historically been regarded as a heterochromatin protein, recruited to pericentromeric foci via the affinity of the ADD domain for H3K9me3 (Dyer et al., 2017). This recruitment is severely disrupted by the R245C mutation, but it is possible that ATRX[R245C] persists at a subset of heterochromatic loci due to recruitment via its interaction partners, such as HP1 and MeCP2. Furthermore, a recent study demonstrated that ATRX is also found at active chromatin sites, such as promoters and enhancers (Truch et al., 2022), where the recruitment mechanism is yet to be determined. ATRX[R245C] retains both the ATPase and DAXX interaction domains, suggesting that if it were able to be recruited to genomic sites, it could remodel chromatin and incorporate histone H3.3.

A patient-relevant mouse model is crucial for therapeutic studies. Our extensive phenotypic characterisation of this model has identified key behavioural paradigms to be used when assessing for phenotypic rescue. The strong phenotypic overlap between the *Atrx^R245C/y^* and *Atrx^Δex2/y^* models supports the relevance of these tests for treating patients carrying all types of ATRX mutations.

## Figures and legends

## Supplementary figures and legends

**Figure S1.**
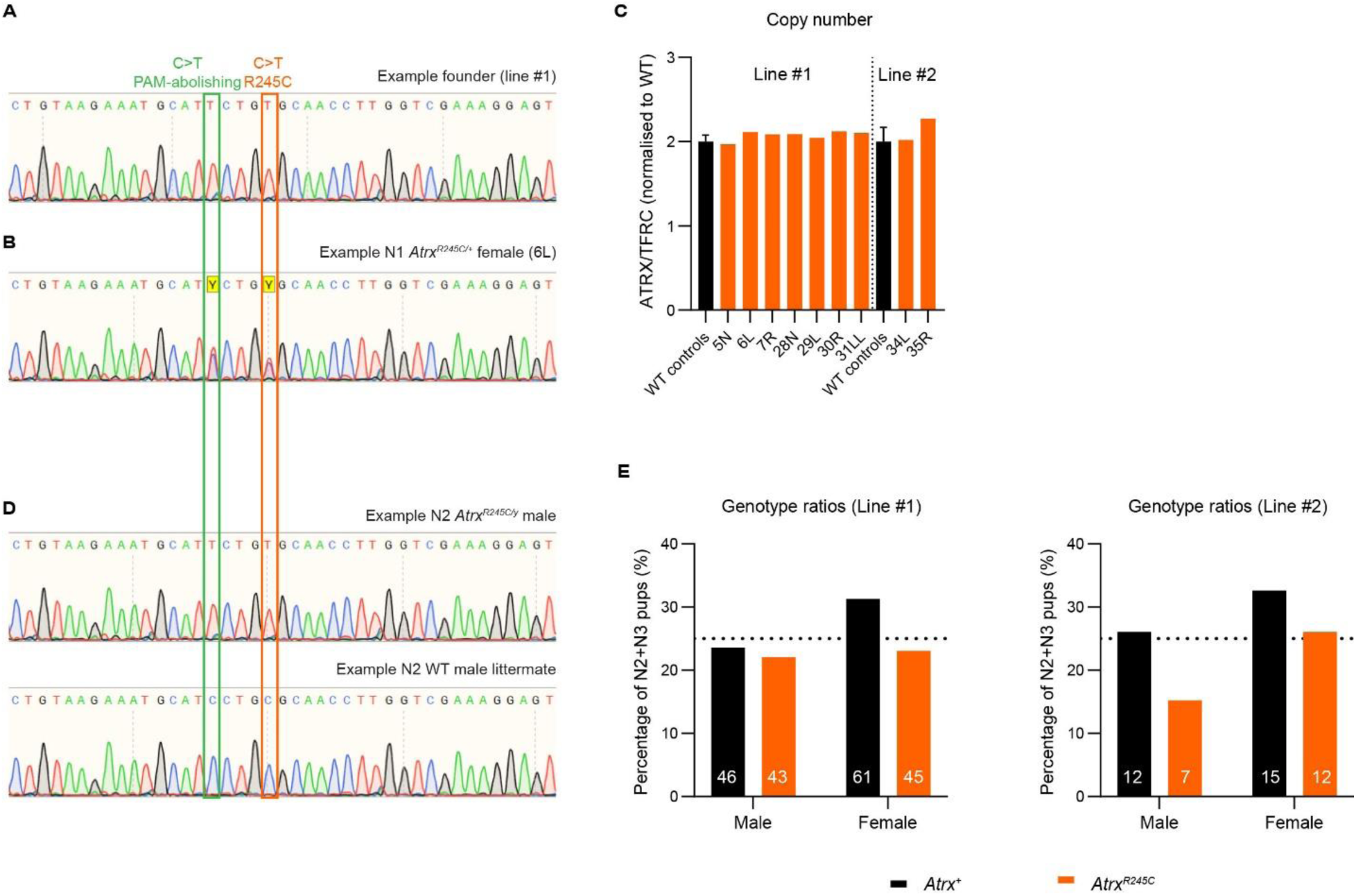
Production of *Atrx^R245C/y^* knock-in mice. **A.** Example Sanger sequencing read from the founder used to establish line #1 (gDNA from tail clip). **B**. Example Sanger sequencing read from an N1 *Atrx^R245C/+^* female, #6L. **C.** qPCR analysis of gDNA using primers within the donor molecule indicate no additional insertions in N1 *Atrx^R245C/+^* females in lines #1 and 2. **D.** Example Sanger sequencing reads from an N2 *Atrx^R245C/y^* male (upper) and WT littermate (lower). **E.** Genotype ratios (%) of N2 and N3 pups in lines #1 and 2 (n numbers on bars). Ratios are not significantly different from expected Mendelian ratios: line #1 P = 0.47; line #2 P = 0.76 (Fisher’s exact tests).

**Figure S2.**
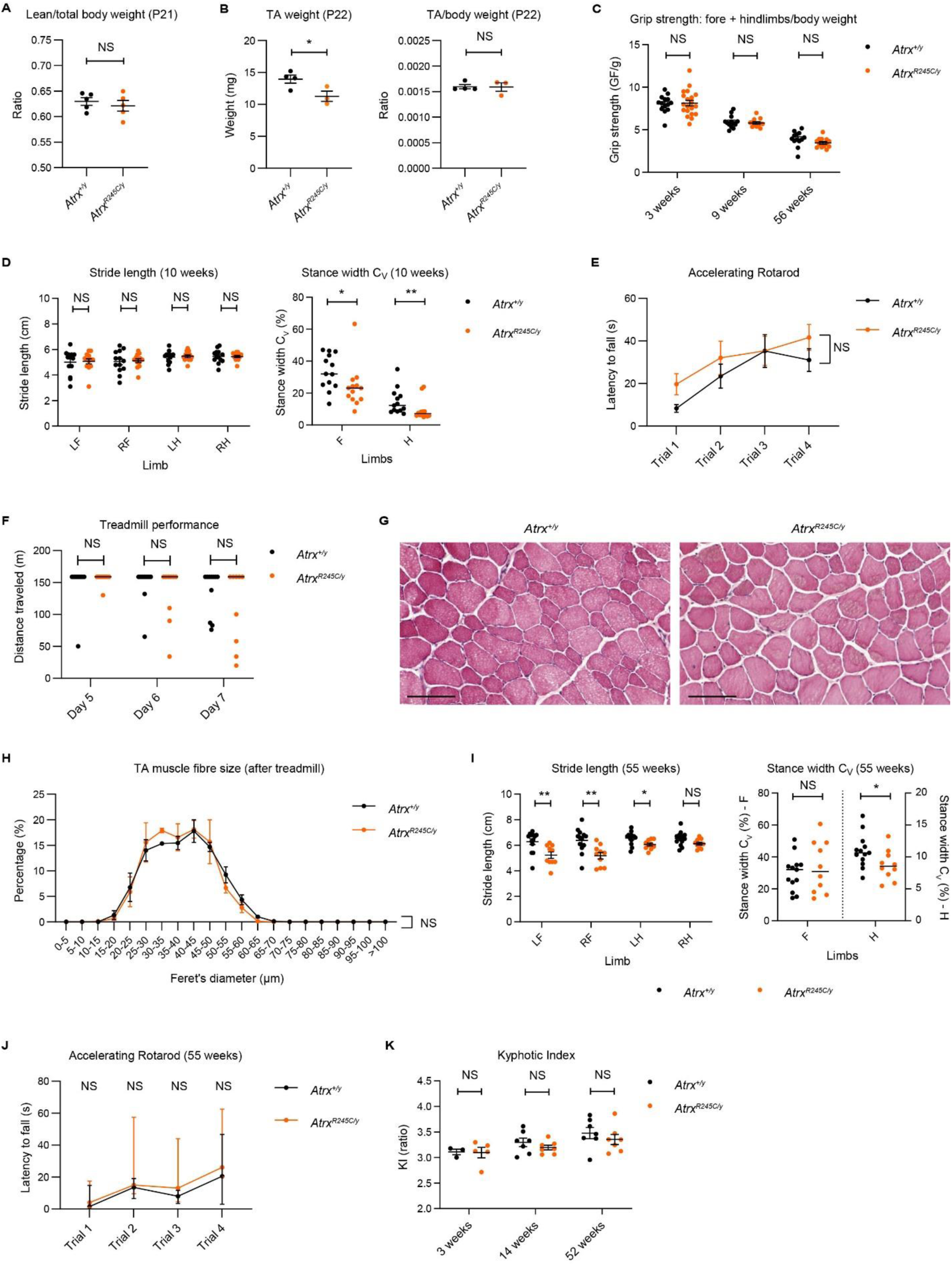
*Atrx^R245C/y^* mice do not display muscle defects. **A.** DEXA analysis of lean/total body weight ratios at P21 (WT n = 5; *R245C* n = 5). Graph shows mean ± S.E.M. and genotypes were compared using a t-test: P > 0.05. **B.** Tibialis anterior (TA) muscle weights (right) and normalised to body weight (left) at P25 (WT n = 4; *R245C* n = 3). Graphs show mean ± S.E.M. and genotypes were compared using t-tests: TA weight: * P = 0.046; TA/body weight P > 0.05. **C.** Forelimb and hindlimb grip strength (normalised to body weight) at 3 weeks (WT n = 14; *R245C* n = 20), 9 weeks (WT n = 13; *R245C* n = 12), and 52 weeks (WT n = 13; *R245C* n = 13). Graph shows mean ± S.E.M. and genotypes were compared using t-tests for each cohort: all P > 0.05. **D.** Gait assessed using motorised treadmill at 19 cm/s: stride length each paw (left) and stance width coefficient of variation (C_V_) for front and hind paws (right) at 10 weeks (WT n = 13; *R245C* n = 13). Stride length: graph shows mean ± S.E.M. and genotypes were compared using t-tests: NS P > 0.05. Stance width C_V_: graph shows medians and genotypes were compared using KS tests: Fore (F) * P = 0.046; Hind (H) ** P = 0.004. **E.** Latency to fall from the accelerating rotarod in four trials over one day at 13 weeks (WT n = 13; *R245C* n = 13). Graph shows mean ± S.E.M. and genotypes were compared by repeated measures ANOVA: NS P > 0.05. Both genotypes show improvement over the course of the day, analysed by 1-way ANOVA: WT ** P = 0.003; R245C: ** P = 0.003. **F.** Performance on treadmill during trials (days 5-7) at 13 weeks (WT n = 11; *R245C* n = 9). Trials were ended early if animals showed signs of exhaustion. Graph shows median and genotypes were compared using Mann-Whitney tests: all P > 0.05. **G.** Representative images of WT (left) and RC (right) tibialis anterior (TA) muscle harvested after chronic exercise, stained with H&E. Scale bar: 100 µm. **H.** Histogram of Feret’s diameter measured in TA muscle after chronic exercise (WT n = 3; *R245C* n = 3). Graph shows mean ± S.E.M. and genotypes were compared using repeated measures ANOVA: P > 0.05. **I.** Gait analysis (as D) of an independent cohort 55 weeks (WT n = 13; *R245C* n = 10). Stride length: graph shows mean ± S.E.M. and genotypes were compared using t-tests: Left fore (LF) ** P = 0.005; Right fore (RF) ** P 0.004; Left hind (LH) * P = 0.015; Right hind (RH) P = 0.055. Stance width C_V_: graph shows medians and genotypes were compared using t-tests: Fore (F) P > 0.05; Hind (H) * P = 0.045. Note: 3/13 mutants were unable to complete the task so were excluded. **J**. Accelerating rotarod analysis (as **E**) of an independent cohort 55 weeks (WT n = 8; *R245C* n = 9). Graph shows median ± interquartile range and genotypes were compared by KS tests: all P > 0.05. Both genotypes show improvement over the course of the day, analysed by Freidman test: WT ** P = 0.009; R245C: ** P = 0.001. **K**. Kyphotic indices measures from X-rays at 3 weeks (WT n = 3; *R245C* n = 5), and 14 and 52 weeks (WT n = 7; *R245C* n = 7). Graph shows mean ± S.E.M. and genotypes were compared using t-tests for each time point: all P > 0.05.

**Figure S3:**
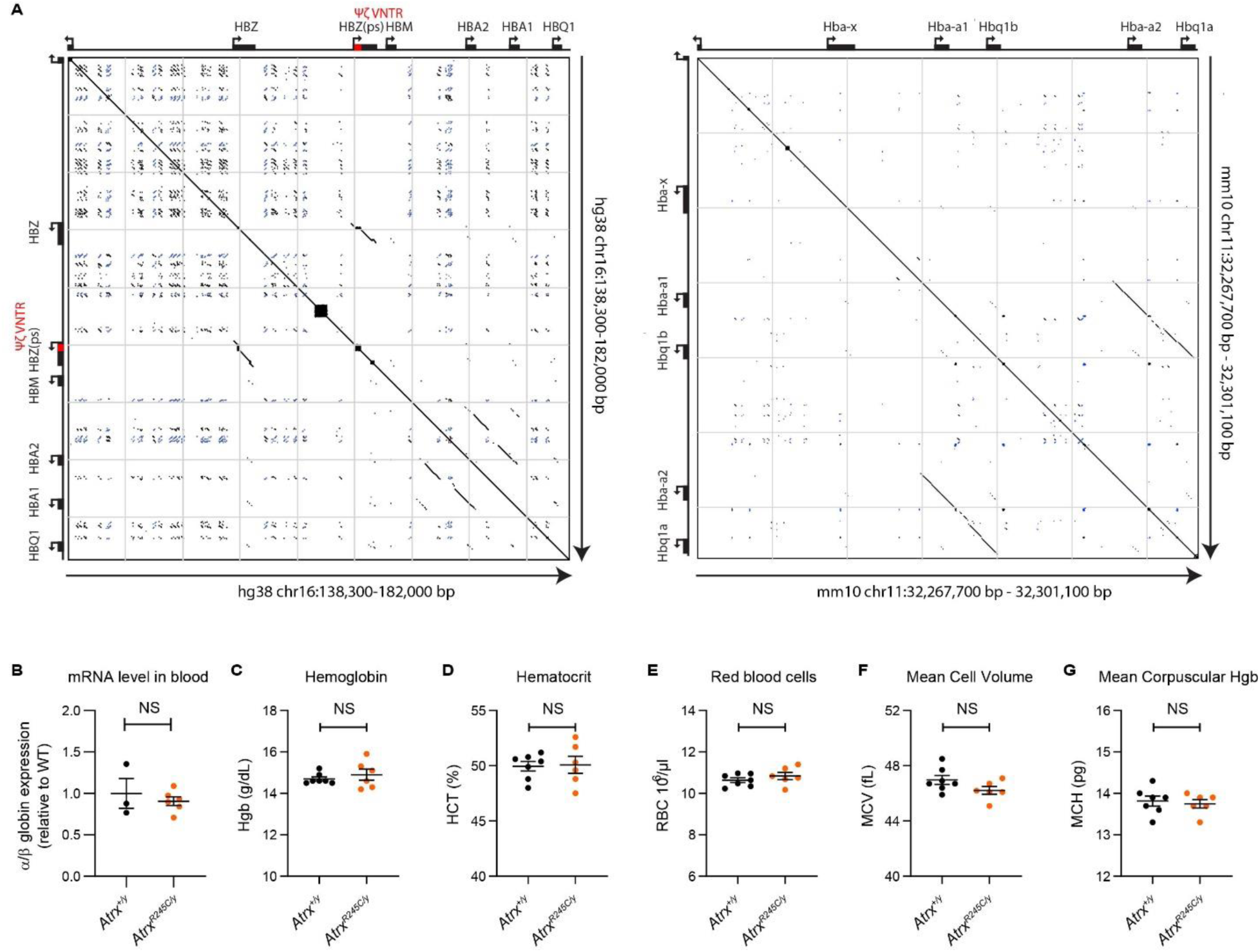
*Atrx^R245C/y^* mice do not develop alpha-thalassemia. **A.** Analysis of self-similarity in the human (left) and mice (right) alpha-globin loci. Dot plots showing self-alignment of a ∼44 kb region from human chromosome 16 and a ∼33 kb region from mouse chromosome 11. Both regions show the region from the NPRL3 promoter to 3’ of the distal paralogous alpha-globin gene. Diagonals represent regions of sequence identity (self-identity diagonal is shown centrally). Black diagonals show identity in the forward orientation and blue diagonals show identity in the reverse orientation. **B.** qPCR analysis of alpha/beta globin expression in blood from *Atrx^R245C/y^* mice and WT controls at 18 weeks (WT n = 3; *R245C* n = 6). Graph shows mean ± S.E.M. and genotypes were compared by a t-test: NS P > 0.05. **C-G.** Complete blood count analysis at 14 weeks (WT n = 7; *R245C* n = 6). Graphs show mean ± S.E.M. and genotypes were compared using t-tests: all not significant (P > 0.05).

**Figure S4.**
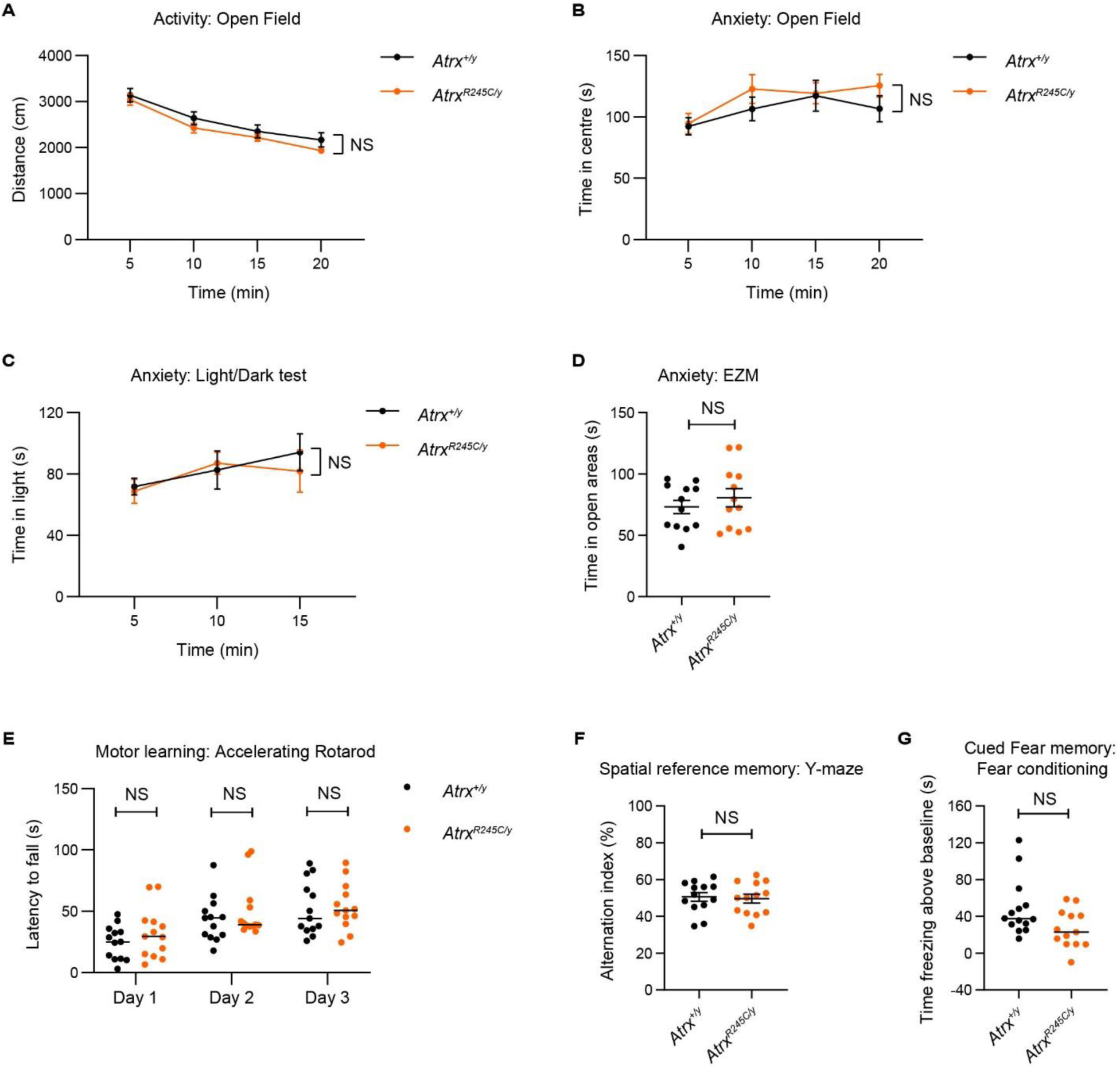
*Atrx^R245C/y^* mice display normal activity, anxiety, motor learning, spatial reference memory and cued fear memory. **A.** Distance travelled over 20 min (as 5 min bins) in the Open Field test at 9 weeks (WT n = 12; *R245C* n = 13). Graph shows mean ± S.E.M. and genotypes were compared using repeated measures ANOVA: P > **B.** Time spent in the centre (40% of total area) in the Open Field test: P > 0.05. **C.** Time spent in the light half of the light/dark test over 15 min (as 5 min bins) at 10 weeks (WT n = 13; *R245C* n = 13). Graph shows mean ± S.E.M. and genotypes were compared using repeated measures ANOVA: P > 0.05. **D.** Time spent in open areas of the elevated zero maze (EZM) over 5 min at 9 weeks (WT n = 12; *R245C* n = 12). Graph shows mean ± S.E.M. and genotypes were compared using a t-test: P > 0.05. **E.** Performance over three days on the accelerating rotarod at 13 weeks (continued from Figure S2E). Means of four daily trials is shown per animal and the line denotes group median (WT n = 13; *R245C* n = 13). Genotypes were compared on each day using KS tests: NS P > 0.05. Both genotypes show learning over the three days of the experiment, analysed by Freidman tests: WT *** P = 0.0008; R245C: * P = 0.037. **F.** Spatial reference memory was assessed over 8 min in the Y-maze test at 12 weeks (WT n = 13; *R245C* n = 13). Alternation index = number of alternations/max alternations * 100. Graph shows mean ± S.E.M. and genotypes were compared using a t-test: P > 0.05. **G.** Cued fear conditioning analysis at 13 weeks (WT n = 14; *R245C* n = 13). Time spent freezing after hearing a tone (minus baseline = before tone) that accompanied the foot shock 24 h previously. Graph shows medians and genotypes were compared using a KS test: P > 0.05.

**Figure S5.**
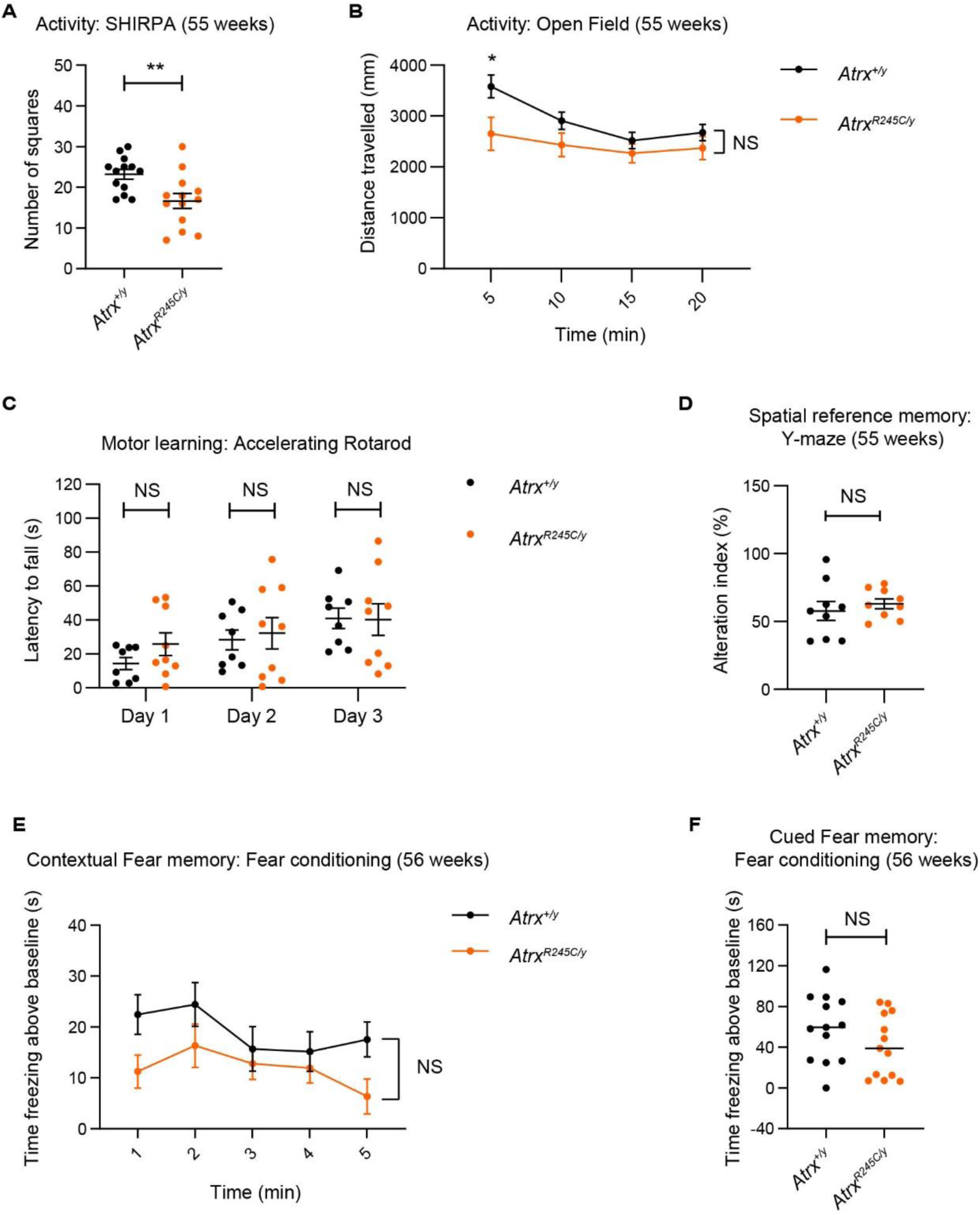
*Atrx^R245C/y^* mice do not develop late onset memory defects. **A.** Spontaneous activity assessed by number of squares entered in 30 s repeated at 55 weeks (WT n = 13; *R245C* n = 13). Graphs show mean ± S.E.M. and genotypes were compared using a t-test: ** P = **B.** Distance travelled over 20 min (as 5 min bins) in the Open Field test at 55 weeks (WT n = 9; *R245C* n = 9). Graph shows mean ± S.E.M. and genotypes were compared using repeated measures ANOVA: P > 0.05. Individual time bins were compared by t-tests and mutants show hypoactivity in bin 1: * P = 0.031. **C.** Performance over three days on the accelerating rotarod at 55 weeks (continued from Figure S2G). Means of four daily trials is shown per animal and the line denotes group mean ± S.E.M. (WT n = 8; *R245C* n = 9). Genotypes were compared on each day using t-tests: NS P > 0.05. Only WT animals show learning over the three days of the experiment, analysed by 1-way ANOVA: WT ** P = 0.006; *R245C* P > 0.05. **D.** Spatial reference memory was assessed over 8 min in the Y-maze test at 55 weeks (WT n = 9; *R245C* n = 9). Alternation index = number of alternations/max alternations * 100. Graph shows mean ± S.E.M. and genotypes were compared using a t-test: P > 0.05. **E-F.** Fear conditioning analysis at 56 weeks (WT n = 13; *R245C* n = 13). **E.** Contextual: time spent freezing (minus baseline determined before shock) when returned to the same environment 24 h after receiving a foot shock. Graph shows mean ± S.E.M. and genotypes were compared using repeated measures ANOVA: P > 0.05. **F.** Cued: time spent freezing after hearing a tone (minus baseline = before tone) that accompanied the foot shock 24 h previously. Graph shows medians and genotypes were compared using a t-test: P > 0.05.

**Figure S6.**
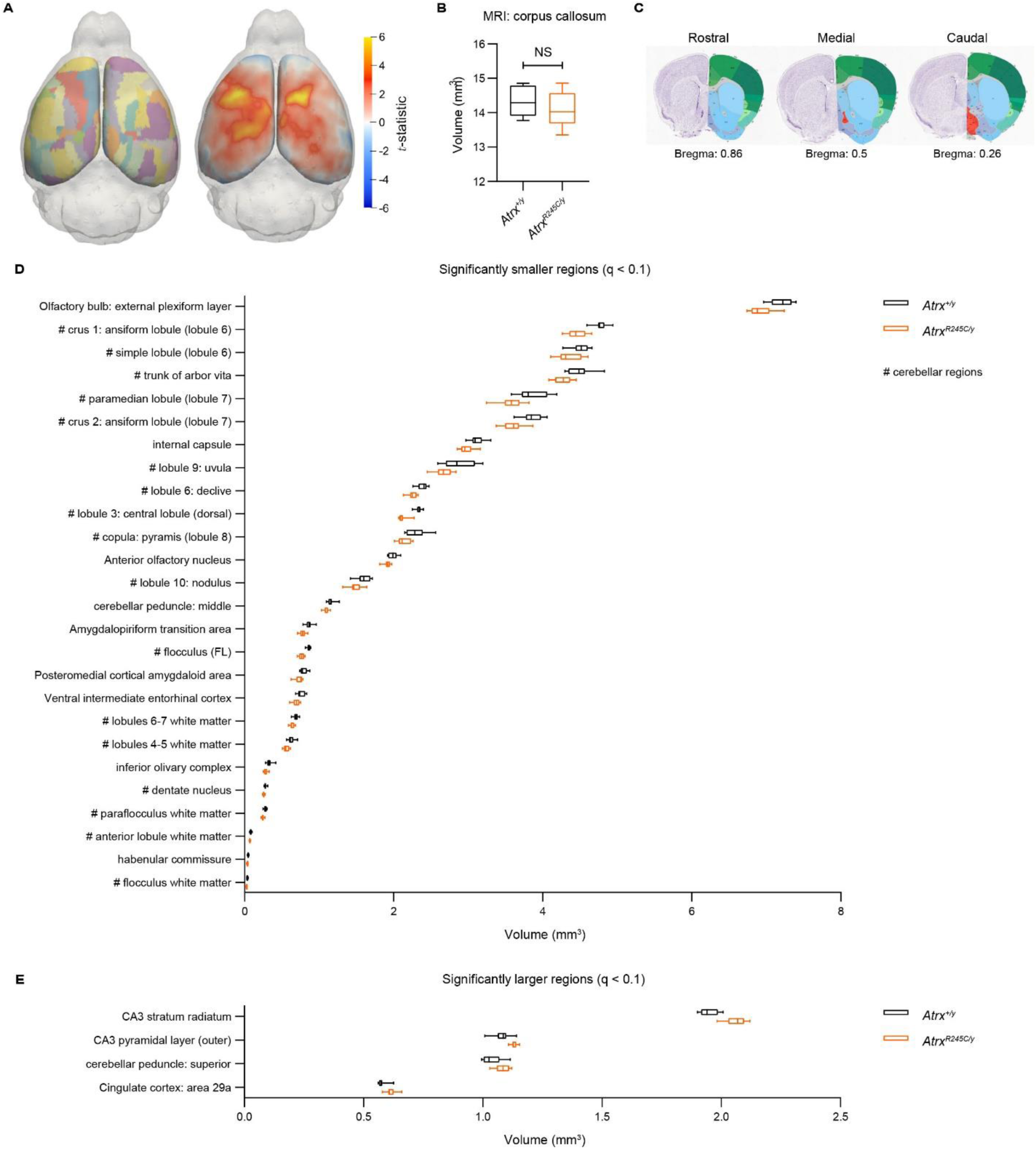
Changes in brain volume in *Atrx^R245C/y^* mice. **A.** Cortical thickness analysed by MRI at 9 weeks (WT n = 10; *R245C* n = 12). Top view showing a surface projection of the DSURQE atlas (left) and cortical thickness differences comparing R245C to WT, computed independently at each surface vertex and quantified by the t-statistic. **B.** Corpus callosum volume. Graph shows interquartile ranges and genotypes were compared using a t-test: P > 0.05. **C.** Diagram showing rostral, redial and caudal sections used to measure corpus callosum thickness. **D-E.** Volumes of the significantly (q < 0.1) smaller (**D**) and larger (**E**) brain regions. Graph shows interquartile ranges with min-max whiskers.

**Figure S7.**
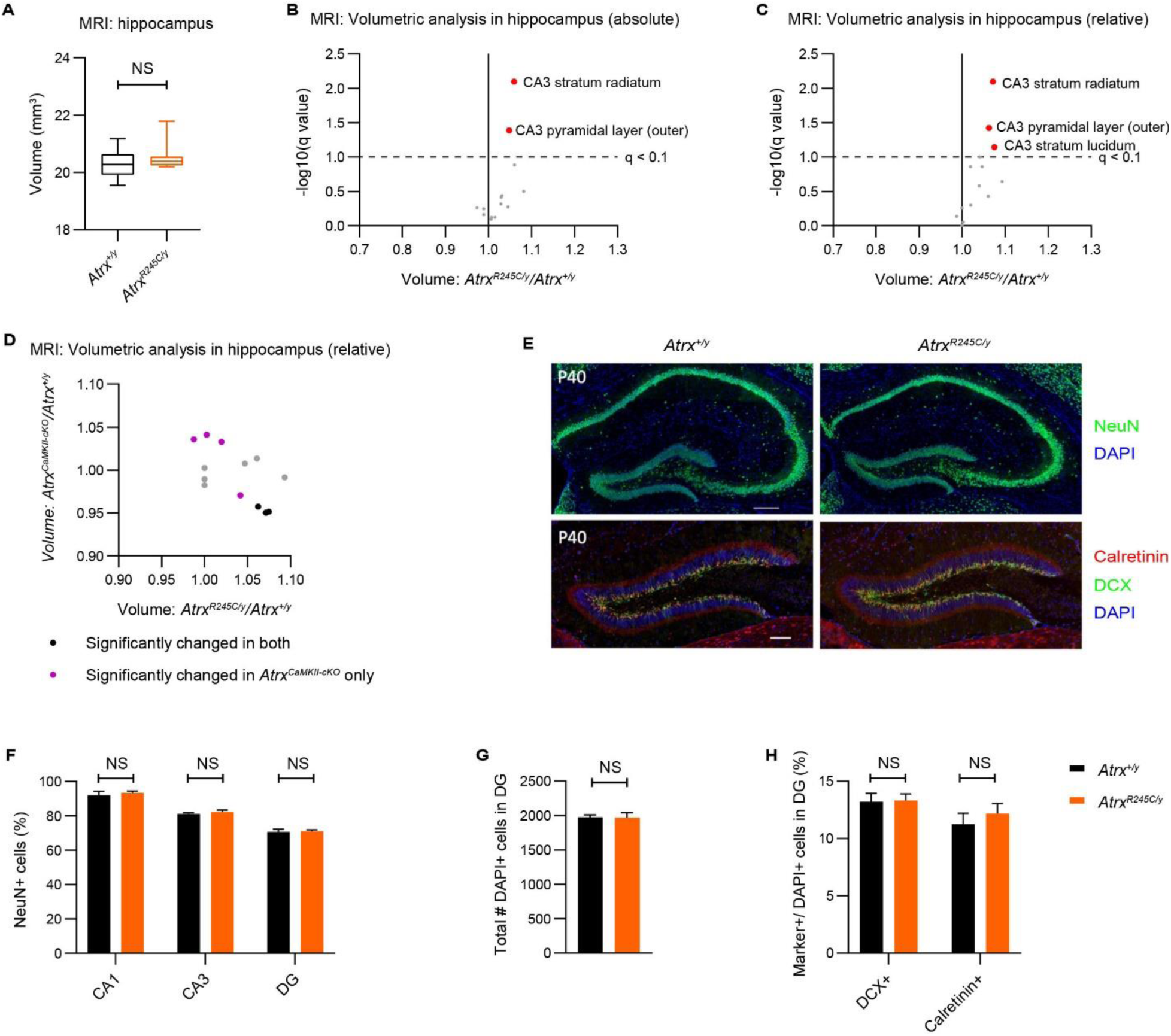
Hippocampal development is unaffected in *Atrx^R245C/y^* mice. **A.** Total hippocampal volume analysed by MRI at 9 weeks (WT n = 10; *R245C* n = 12). Graph shows interquartile ranges and genotypes were compared using a KS test: P > 0.05. **B-C.** Volcano plots of absolute (**B**; as in Figure 6C,E) and relative (**C**; as performed for *Atrx CaMKII-cKO* mice (Tamming et al., 2020)) volumetric analysis of the 15 subregions of the hippocampus. Significantly increased (q < 0.1) regions are highlighted in red. **D.** Comparison of relative volumetric analysis in the hippocampus between *Atrx^R245C/y^* and *Atrx CaMKII-cKO* mice. **E.** Representative images of hippocampal (upper) and dentate gyrus (lower) sections from WT and R245C mice at P40. Hippocampal sections were stained with NeuN a marker of mature neurons (green). Scale bar: 200 µm. Dentate gyrus sections were stained with calretinin (red), DCX (green) and DAPI. Scale bar: 100 µm. **F.** Percentage of NeuN+ cells for CA1, CA3 and dentate gyrus (DG) regions of the hippocampus (see Fig S9E, upper). **G-H.** Total number of DAPI+ cells (**G**) and number of calretinin+ and DCX+ cells (**H**) in the DG (see Figure S9E, lower). **E-H.** N = 4 biological replicates per genotype. All graphs show mean ± S.E.M. and genotypes were compared using t-tests: all NS P > 0.05.

**Table S1:**
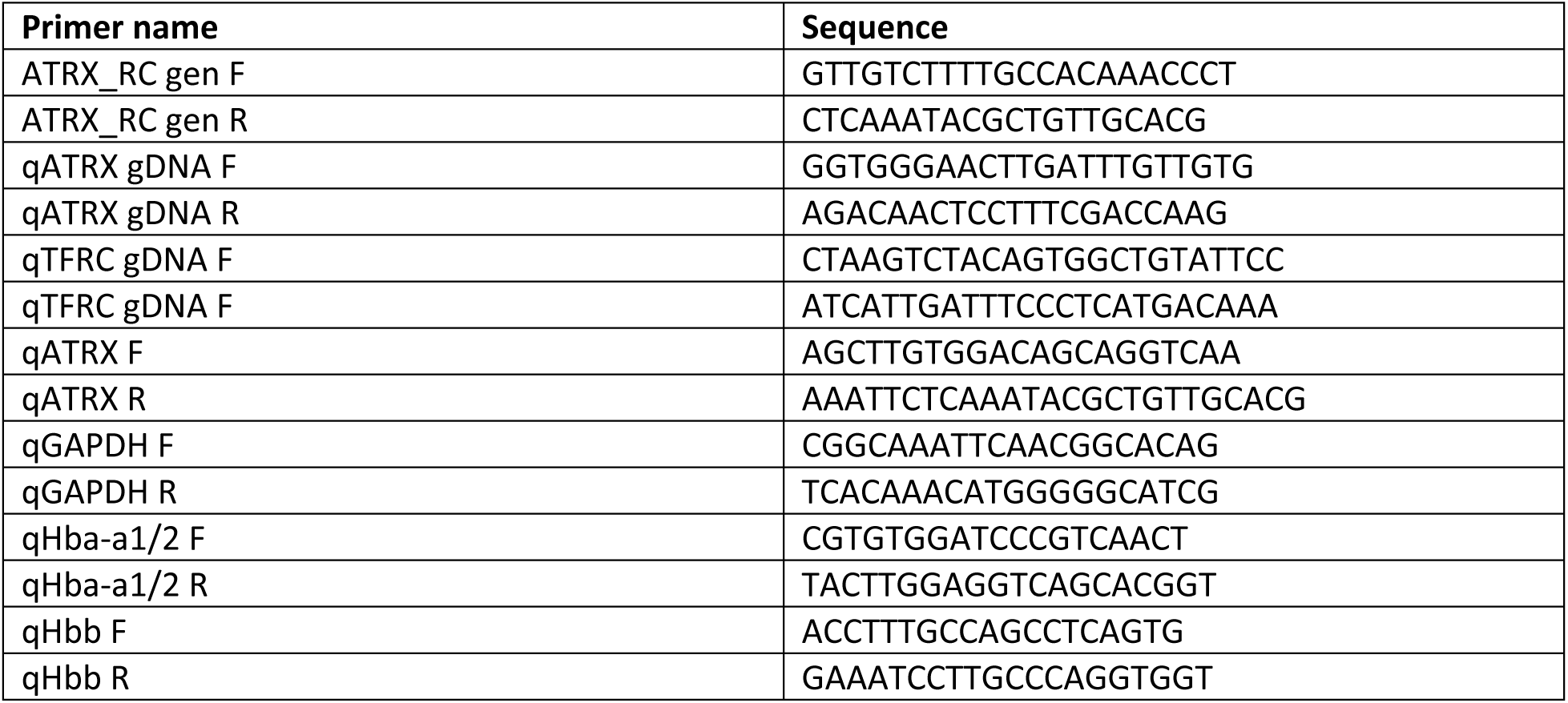
Primers

## Acknowledgements

The authors wish to acknowledge the contribution of Lauryl Nutter and Marina Gertsenstein in The Model Production Core at The Centre for Phenogenomics for the generation of the *Atrx^em1Tcp^* mutant mice and Ann Flenniken and Zorana Berberovic in in the Phenotyping Core at The Centre for Phenogenomics for X-ray imaging, DEXA and blood analysis. We thank Igor Vukobradovic, Gabi Gurria, Sebastian Gerety and Matthew Hurles for advice on behavioural testing and analysis of craniofacial morphology; Eleonora Maino for advice on muscle analysis; and Tiffany Chien, Ron Padilla and Zachary Klugman for technical assistance. We also thank members of the Justice, Picketts, R. J. Gibbons and D. R. Higgs laboratories for helpful discussions.

**Conflict of interest statement:** the authors have no conflicts of interest to declare.

## Author contributions

Conceptualization, R.T., M.J.J., and D.J.P.; methodology, R.T., K.Y., T.Y., and A.C.; software, T.Y., J.G.S and B.J.N; formal analysis, R.T., T.Y., K.Y., A.C., V.T.-C., Y.L. and C.B.; investigation, R.T., K.Y., A.C. V.T.-C., C.T., S.V. and J.R.; writing – original draft, R.T.; writing – review & editing, all of the authors; visualization, R.T., K.Y., T.Y. A.C., V.T.-C. and Y.L.; supervision, M.J.J., D.J.P., J.G.S., B.J.N., and E.A.I.; project management, J.R; funding acquisition, R.T., M.J.J., and D.J.P..

## Materials and Methods

### Experimental model and subject details

#### Mouse lines

Animal procedures were approved by the Animal Use Committees at the Canadian Council on Animal Care (CCAC)-accredited animal facilities, The Center for Phenogenomics (TCP) and the University of Ottawa. All mice were housed in specific-pathogen-free (SPF) facilities. They were maintained on a 12-h light/dark cycle and given ad libitum access to food and water. Mice were fed a standard diet (HarlanTeklad 2918) ad libitum, consisting of 18% protein, 6% fat and 44% carbohydrates. They were housed in individually ventilated cages (IVC) with wood chippings, tissue bedding and additional environmental enrichment in groups of up to five animals. Mutant mice were housed with their wild-type littermates. *Atrx^em1Tcp^* (*referred to as Atrx^R245C^*) mice were generated in this study the TCP model production core via CRISPR/Cas9-meditated editing in C57BL/6J zygotes (detailed method described below). Heterozygous *Atrx^R245C/+^* females were crossed with wild-type C57BL/6J males to produce hemizygous *Atrx^R245/y^* males and wild-type littermate controls for all experiments.

Cohort 1 (*Atrx^+/y^* n = 13; *Atrx^R245C/y^* n = 13) underwent body weight measurements (5-52 weeks), SHIRPA (9 weeks), Social Interaction (10 weeks), Y-maze (11 weeks), Accelerating Rotarod (13 weeks), SHIRPA (55 weeks), gait analysis (55 weeks), grip strength measurement (55 weeks), Fear Conditioning (56 weeks). Cohort 2 (*Atrx^+/y^* n = 14; *Atrx^R245C/y^* n = 13) underwent Open Field (9 weeks), Light/Dark box (10 weeks), gait analysis (10 weeks), Barnes Maze (12 weeks) and Fear Conditioning (13 weeks) and complete blood counts (14 weeks). Cohort 3 (*Atrx^+/y^* n = 11; *Atrx^R245C/y^* n = 9) underwent grip strength measurement (3 weeks) and chronic exercise (13 weeks). Cohort 4 (*Atrx^+/y^* n = 13; *Atrx^R245C/y^* n = 12) underwent Elevated Zero Maze (9 weeks) and grip strength (9 weeks) and MRI (9 weeks). Cohort 5 (*Atrx^+/y^* n = 9; *Atrx^R245C/y^* n = 9) underwent Open Field (55 weeks), Y-maze (55 weeks) and Accelerating Rotarod (55 weeks). Cohort 6 (*Atrx^+/y^* n = 5; *Atrx^R245C/y^* n = 5) underwent DEXA (3 weeks). Cohort 7 (*Atrx^+/y^* n = 3; *Atrx^R245C/y^* n = 11) underwent grip strength measurement (3 weeks) and X-ray analysis (3 weeks; *Atrx^+/y^* n = 3; *Atrx^R245C/y^* n = 5). Cohort 8 (*Atrx^+/y^* n = 7; *Atrx^R245C/y^* n = 7) underwent X-ray analysis (14 and 52 weeks). Cohort 9 (*Atrx^+/y^* n = 7; *Atrx^R245C/y^* n = 8) underwent X-ray analysis (40 weeks).

### Method details

#### Generation of Atrx^em1Tcp^ (referred to as Atrx^R245C^) knock-in mice

*Atrx^em1Tcp^* mice were generated by the Model Production Core at The Centre for Phenogenomics by injecting Cas9 endonuclease and a guide RNA with the spacer sequence CCTTTCGACCAAGGTTGCGC and a single-strand oligonucleotide encoding the changes c.733C>T, p.R245C and c.729C>T, p.I243I to inactivate the PAM sequence in ENSMUST00000113573 (NM_009530) encoding NP_033556. These reagents were introduced into C57BL/6J zygotes (Gertsenstein and Nutter, 2021) and maintained on this background by crossing heterozygous *Atrx^R245C/+^* females with wild-type C57BL/6J males.

Introduction of the desired mutations was assessed in founders by Sanger sequencing after PCR amplification. This was confirmed in N1 heterozygous *Atrx^R245C/+^* females after digestion of the wild-type allele using FspI (NEB R0135S) and gel purification of the uncut mutant band. Copy number was analysed by qPCR amplification of PCI-purified genomic DNA (diluted to 2.5 ng/µl) to verify that no additional copies of the donor oligonucleotide had integrated. The ATRX locus was detected with qATRX F and qATRX R primers (which lie within the donor oligonucleotide). Values were normalised to the TFRC locus, amplified with qTFRC F and qTFRC R primers. qPCR was performed on a Viia7 instrument (ABI) instrument using SYBR Green PCR Master Mix (Invitrogen). PCR amplification conditions: annealing temperature 58°C, 40 cycles. Relative abundance = E^TFRC^^CT^TFRC^ / E^ATRX^^CT^ATRX^. Abundance was normalised to the mean of three heterozygous *Atrx^R245C/+^* females. See Table S1 for primer sequences. Genotype ratios of N2 and N3 pups were analysed using Fisher’s exact tests.

#### Genotyping

Tail tip genomic DNA was PCR amplified with ATRX_RC gen F and ATRX_RC gen R primers and digested with FspI (NEB R0135S). FspI cuts the wild-type allele (product sizes: 245 + 120 bp), but the restriction site is lost in the R245C allele (product size: 365 bp). See Table S1 for primer sequences.

#### Neuronal cell cultures

Cortical and hippocampal neuron cultures were prepared from E17.5 mouse embryos, based on Wilson et al., 2020. Briefly, embryos were individually decapitated and both cortex or hippocampi dissected and collected in cold Hank’s balanced salt solution (HBSS: NaHCO3 4.2 mM, Hank’s salt powder 0.952%, HEPES 12 mM, 4-(2-hydroxyethyl)-1-piperazineethane-sulphonic acid, Sigma), then digested with 0.25% Trypsin at 37°C for 8 minutes. Enzymatic digestion was blocked with Dulbecco’s Modified Eagle Medium high glucose (DMEM, supplemented with 10% Fetal Bovine Serum (FBS) and penicillin-streptomycin). The tissue was centrifuged at 800 rpm for 5 minutes at room temperature. The tissue was resuspended with DMEM + 10% FBS and mechanically triturated. Hippocampal neurons were plated on 12 mm cover glasses and cortical neurons directly on polystyrene multiwell plates, coated with 1mg/ml poly-L-lysine (Sigma). Cultures were grown at 37°C and 5% CO_2_ in Neurobasal (Invitrogen) supplemented with 2% B-27 (Invitrogen), 1 mM L-glutamine and 1% penicillin-streptomycin. Genotype and sex were later confirmed by PCR.

#### RNA purification and qPCR

Cortices were dissected from *Atrx^R245C/y^* mice and *Atrx^+/y^* littermate controls at P0.5 and P60 RNA was isolated using Trizol (Thermo Fisher Scientific #15596026) according to the manufacturer’s instructions. 2 µg of total RNA was used for DNase treatment using DNA-free™ DNA Removal Kit (Fisher #AM1906) and then reverse-transcribed to cDNA using RevertAid Reverse Transcriptase (Fisher #EP0442) according to the manufacturer’s instructions. Synthesized cDNA was further diluted 1:10 for qPCR. RT-qPCR analysis was carried out using the Lo-Rox SYBR 2x mix (FroggaBio Inc #CSA-01195) under the following conditions: one cycle at 95°C for 1 min, and then 40 consecutive cycles at 95°C for 10 s, 60°C for 10 s, and 72°C for 20 s using AriaMx Real-Time PCR System (Agilent). All primers (Table S1) were analyzed by melt curve analysis after qPCR amplification. The ΔΔCt method was used to compare fold-change. GAPDH was for normalisation. Triplicate or quadruplicate samples were performed per reaction and a minimum of 3 mice analyzed per genotype. Student’s t-test was used for statistical significance.

Whole blood was collected from the saphenous vein in EDTA at 18 weeks (*Atrx^+/y^* n = 3; *Atrx^R245C/y^* n = 6). 100 µl whole blood was homogenised in 1 ml QIAzol (Qiagen) using a TissueLyser II (Qiagen) at 20 Hz for 3 min. Samples were incubated at room temperature for 5 min before 200 µl chloroform was added. Samples were centrifuged at 12,000 g for 15 min at 4°C and the upper phase was transferred to a new tube. RNA was precipitated with 600 µl 70% ethanol and isolated in a RNeasy Mini Spin Column. RNA was purified using the RNeasy kit (Qiagen 74106) according to manufacturer’s instructions. In short, the RNA was washed with 700 µl Buffer RW1, then 2x 500 µl Buffer RPE, and eluted in 50 µl RNAse-free water. gDNA removal and cDNA synthesis was performed from 1.25 µg RNA using SuperScript™ IV VILO™ Master Mix with ezDNase™ Enzyme (Thermo Scientific 11766050) according to manufacturer’s instructions. Diluted cDNA 1 in 1000 for analysis of α-globin expression by qPCR amplified with qHba-a1/2 F and qHba-a1/2 R, normalised to β-globin expression amplified with qHbb F and qHbb R. See Table S1 for primer sequences. PCR amplification conditions: annealing temperature 60°C, 40 cycles. Relative abundance = E^Hbb^^CT^Hbb^ / E^Hba^^CT^Hba^. Genotypes were compared using t-tests (unpaired, two-tailed).

#### Protein purification and Western blotting

Cortices were quickly dissected, snap frozen in liquid nitrogen, and then homogenized in ice cold RIPA buffer (10 mM Tris-Cl, pH8.0, 1 mM EDTA, 1% Triton X-100, 0.1% sodium deoxychlorate, 0.1% SDS, 140 mM NaCl, and 1 mM PMSF) supplemented with protease inhibitor cocktail (Sigma-Aldrich). After 20 min incubation on ice, samples were centrifuged (17,000g for 20 min) and the protein supernatant quantified by Bradford assay. Protein samples were resolved on sodium dodecyl sulfate polyacrylamide gels under denaturing conditions and blotted onto PVDF membranes (Immobilon-P; Millipore, Burlington, MA, United States) by wet transfer for 1–2 h at 90 V. Membranes were blocked (45 min, room temperature) with 5% skim milk in TBST (Tris-buffered saline containing 0.05% Triton X-100), and incubated (4°C, overnight) with the following antibodies: mouse anti-ATRX 39F (1: 1000); rabbit anti-Tbr2 (1:1000; Abcam #37003); mouse anti-Satb2 (1:500; Abcam #51502); rat anti-Ctip2 (1:2000; Abcam #18465); rabbit anti-Tbr1 (1:1000; Abcam #31940; mouse anti-ß-Actin (1:30,000; Sigma #A5441); and mouse anti-Vinculin (1:10,000; Sigma-Aldrich #V9131). Membranes were incubated (1 h, room temperature) with HRP-conjugated goat anti-Rabbit IgG (Sigma #A4914), sheep anti-mouse (Sigma #A5906) or goat anti-rat (Sigma #AP136P) secondary antibodies (1:3000 to 1:30,000). Membranes were washed 5 X 5 min in TBST and signals were detected using the Clarity Western ECL Substrate (Biorad #1705060). At least 2 separate gels were immunoblotted with cortical extracts from independent litters and used for quantitation.

#### Cyclohexamide assay

For analysis of protein stability, cultures were treated with vehicle (DMSO) or 100 µM CHX (Sigma) for 0, 3, and 8 h. Cells were lysed in RIPA lysis buffer and protein was collected for western.

#### Nissl staining and Immunofluorescence

Nissl staining was performed using cresyl violet standard staining procedures. In brief, 12 µm frozen sections were rehydrated in 95%, 70%, and 50% ETOH (ethanol) for 10 min, 1 min, and 1 min, respectively, and then incubated in ddH2O for 2 X 5 min. Rehydrated sections were then stained in 1% cresyl violet acetate staining solution (1% cresyl violet, 0.25% glacial acetic acid) for 4 min. After 5 min washing in ddH2O, the sections were dehydrated in 50%, 70%, 95% and 100% ETOH for 2 min, 2 min, 2 min, and 5 min, respectively. Sections were then cleared by incubation in xylene for 3 X 5 min, mounted with Permount and visualized under Axio Scan. Z1 (Zeiss) scanning microscope.

Postnatal day 0.5 (P0.5) brain sections (12um) were washed 3 X 5 min in PBST (PBS with 0.1% Triton X-100) prior to blocking or antigen retrieval. For antigen retrieval, slides were submerged in citrate buffer (recipe) and heated in a microwave (power level 2) for 10 min. Slides were blocked for 1 hour at room temperature in 10% horse serum/PBST, and then incubated (overnight, 4°C) with primary antibodies. The following primary antibodies were used at 1:200, unless indicated: rabbit anti-Tbr2 (Abcam #37003); rabbit anti-Pax6 (1:100; Abcam #195045); mouse anti-SATB2 (1:50; Abcam #51502); rat anti-Ctip2 (Abcam #18465); rabbit anti-Tbr1 (Abcam #31940); mouse anti-ATRX (39F; (McDowell et al., 1999)). The following day, sections were washed 4 X 5 min in PBST and then incubated with DyLight488, DyLight594, or DyLight649-conjugated secondary antibodies (1:500; Jackson ImmunoResearch, PA). All sections were counterstained with Hoescht 33342 dye (Thermo Fisher Scientific) and coverslips were mounted with Dako Fluorescence Mounting Medium (Dako Canada, ON).

#### Morphological and muscle function analysis

Dual-energy X-ray absorptiometry (DEXA) was performed on anaesthetised mice by The Clinical Phenotyping Core at The Centre for Phenogenomics at 21 days of age (*Atrx^+/y^* n = 5; *Atrx^R245C/y^* n = 5). X-rays were performed on anaesthetised mice (Faxitron MX-20) by The Clinical Phenotyping Core at The Centre for Phenogenomics at 3 weeks (*Atrx^+/y^* n = 3; *Atrx^R245C/y^* n = 5); 14 and 52 weeks (*Atrx^+/y^* n = 7; *Atrx^R245C/y^* n = 7) and 40 weeks (*Atrx^+/y^* n = 7; *Atrx^R245C/y^* n = 8).

Grip strength was measured on a Grip Strength Meter (Bioseb) using three separate cohorts aged 3 weeks (*Atrx^+/y^* n = 14; *Atrx^R245C/y^* n = 20), 9 weeks (*Atrx^+/y^* n = 13; *Atrx^R245C/y^* n = 12) and 52 weeks (*Atrx^+/y^* n = 13; *Atrx^R245C/y^* n = 13). Settings: GF (gram-force); mode = T-PK). Animals were placed with their torso parallel to the grid (no mesh on grid) and allowed to attach before pulling backwards over the grid by its tail. The mean of three measurements for fore + hindlimbs was calculated and normalised by body weight. Gait was analysed on a DigiGait treadmill (Mouse Specifics, Inc.) using two separate cohorts aged 10 weeks (*Atrx^+/y^* n = 13; *Atrx^R245C/y^* n = 13) and 55 weeks (*Atrx^+/y^* n = 13; *Atrx^R245C/y^* n = 10, as 3/13 *Atrx^R245C/y^* failed to walk on the treadmill). 5-8 s of continuous movement were recorded at 19 cm/s (no slope). Videos were analysed using DigiGait Analysis 15 software. A cohort of mice underwent chronic exercise on a treadmill (Columbus Instruments 1055SRM Exer-3/6 Open Treadmill) at 13 weeks (*Atrx^+/y^* n = 11; *Atrx^R245C/y^* n = 9). Mice were acclimatised to the treadmill over four training days: day 1 = 3 metres/min for 1 min, no slope; day 2 = 3 metres/min for 1 min then 4 metres/min for 1 min, 5° decline; day 3 = 3 metres/min for 1 min, 4 metres/min for 1 min then 5 metres/min for 1 min, 10° decline; and day 4 = 3 metres/min for 1 min, 4 metres/min for 1 min, 5 metres/min for 1 min, then 6 metres/min for 3 min, 15° decline; before undergoing trials for three days (days 5-7): each 3 metres/min for 1 min, 4 metres/min for 1 min, 5 metres/min for 1 min, 6 metres/min for 3 min, 8 metres/min for 0.5 min, 10 metres/min for 0.5 min then 12 metres/min for 10 min, 15° decline. Animals were manually kept on the treadmill by the experimenter (shock grids were not used) and monitored for signs of exhaustion, at which point they were returned to their home cage to recover. Mice were injected with Evans Blue Dye (1% w/v in PBS) via IP (10 µl/g body weight) after the second trial day to visualise muscle damage. TA muscle was harvested, coated in OCT and snap-frozen in isopentane chilled in liquid nitrogen. Muscle tissue from the worst three performing animals per genotype (to maximise the chance of identifying damage) was sectioned at 8 µm on a Cryostat for analysis. Infiltration of Evans Blue Dye was analysed by fluorescence microscopy (not detected). Muscle sections were stained with hematoxylin and eosin (H&E) and slides were then scanned using the 3DHPannoramic Slide Scanner by the Imaging Facility at the Hospital for Sick Children. SlideViewer (3DHISTECH) was used for image acquisition. Feret’s diameter was analysed using Image J to quantify MinFeret (n = 300 myofibres per animal).

#### Magnetic resonance imaging (MRI)

MRI for assessment of brain morphology was performed *in vivo* at the Mouse Imaging Centre using multiple-mouse MRI, imaging up to four mice simultaneously (Arbabi et al., 2022; Nieman et al., 2018). Twenty-four hours prior to scanning, mice received an intraperitoneal injection of 30 mM MnCl_2_ (0.4 mmol per kg; M8054, Sigma-Aldrich). During the MRI scan, mice were anesthetized with isoflurane (1-1.5%) and monitored based on their respiratory signal (Nieman et al., 2009). Gradient-echo images were acquired with the following parameters: TR = 26 ms; TE = 8.25 ms; 2 averages; 334×294×294 matrix size; 25×22×22 mm^3^ field of view; 75 μm isotropic voxel size; and a total acquisition time of 1 hr using a “cylindrical” masking of k-space (Spencer Noakes et al., 2017).

#### Haematology

Complete blood counts were performed by The Clinical Phenotyping Core at The Centre for Phenogenomics. Genotypes were compared using t-tests (unpaired, two-tailed).

#### Behavioural analysis

Modified SHIRPA (SmithKline Beecham, Harwell, Imperial College, Royal London Hospital, phenotype assessment) screening was performed on the same cohort at 9 and 55 weeks of age (*Atrx^+/y^* n = 13; *Atrx^R245C/y^* n = 13). Animals were assessed for appearance, spontaneous locomotor activity (number of squares entered with all four paws in a 51 x 37 cm arena marked with 15 equal squares in 30 s), gait, reflexes (startle response, touch escape, righting reflex, contact righting reflex in plastic tube, visual placing, pinnal reflex, corneal reflex, response to tail pinch and pupillary light reflex), behaviour during screen and when handled (passivity, trunk curl, limb grasping, biting and vocalisation. Open Field was performed on two independent cohorts at 9 weeks (*Atrx^+/y^* n = 12; *Atrx^R245C/y^* n = 13) and 55 weeks (*Atrx^+/y^* n = 9; *Atrx^R245C/y^* n = 9). Animals explored an evenly lit (200 lux) 43.5 x 43.5 cm arena for 20 min and their movement was detected using 16 beam IR arrays (X and Y axes). Data was processed using Activity Monitor software. The centre of the area was defined as the central 40% of the total area. The Light/Dark test was performed at 9 weeks (*Atrx^+/y^* n = 13; *Atrx^R245C/y^* n = 13). Lidded dark box inserts were added to the Open Field arena (50% of arena area). Animals were placed in the dark half and allowed to explore for 15 min and their movement was detected using 16 beam IR arrays (X and Y axes). Data was processed using Activity Monitor software. The Elevated Zero Maze was performed at 9 weeks (*Atrx^+/y^* n = 12; *Atrx^R245C/y^* n = 12). Animals explored a “O” shaped maze with diameter 60 cm and path width 7 cm, divided into four equal quadrants: two open quadrants and two walled quadrants, elevated 70 cm off the floor for 5 min. Behaviour was monitored and analysed using Ethovision XT 13 software (Noldus). The three day accelerating rotarod test was performed on two independent cohorts at 13 weeks (*Atrx^+/y^* n = 13; *Atrx^R245C/y^* n = 13) and 55 weeks (*Atrx^+/y^* n = 8; *Atrx^R245C/y^* n = 9). Animals were placed onto a rod facing forwards on the apparatus (Accelerating RotaRod for 5 mice LE8200, Harvard Apparatus Canada). The motor was then started on acceleration mode from 4-40 rpm over 5 min. The latency to fall for each trial was recorded. Each day consisted of four trials, separated by an intertrial interval of 30-45 min. Baseline motor function was assessed on day 1 and motor learning ability was assessed over the total three days. Barnes maze analysis was performed at 12 weeks (*Atrx^+/y^* n = 14; *Atrx^R245C/y^* n = 12) over five consecutive days. The maze consisted of a circular white evenly lit (∼75 lux) platform (121 cm in diameter) elevated ∼1 m from the floor with 40 equally spaced holes around the periphery. A black escape box was places under one hole and shallow black boxes were placed under the remaining holes. The maze was surrounded by four spatial clues to aid orientation. On day 1, mice were placed into a cylindrical holder in the centre of the maze and released after 10 s to explore and find the escape box for a maximum of 3 min. If the subject did not enter the escape box, it was gently guided by the experimenter. Mice were allowed to stay in the escape box for 10 s. On days 2-5, mice underwent four trials (with an intertrial interval of 2 hours), where they were given a maximum of 3 min to enter the escape box. If the subject did not enter the escape box, it was gently guided by the experimenter. Mice were allowed to stay in the escape box for 10 s. For all trials, music was played from when the mouse was released until it entered the escape box. Behaviour was monitored and analysed using Ethovision XT 14 software (Noldus) using three-point tracking. Social novelty was assessed at 10 weeks (*Atrx^+/y^* n = 12; *Atrx^R245C/y^* n = 13) and an arena comprising three equal chambers. Animals were first acclimatised to the empty arena for 5 min. In Session I, two cups were added in opposite corners one of which contained “stranger 1” and animals were released from the central chamber and allowed to explore for 10 min. In Session I, “stranger 1” remained in place and “stranger 2” was added to the second cup and animals were released from the central chamber and allowed to explore for 10 min. All strangers were male C57BL/6J mice around 10 weeks of age. Behaviour was monitored and analysed using Ethovision XT 13 software (Noldus) using three-point tracking. Y-maze was performed on two independent cohorts at 11 weeks (*Atrx^+/y^* n = 13; *Atrx^R245C/y^* n = 13) and 55 weeks (*Atrx^+/y^* n = 9; *Atrx^R245C/y^* n = 9). The Y-shaped arena consists of three arms of identical length at 120° angles. Mice were placed at the end of one arm (facing the end) and allowed to explore for 8 min. Behaviour was monitored and analysed using Ethovision XT 13 software (Noldus). Contextual and cued fear conditioning was performed on two independent cohorts at 13 weeks (*Atrx^+/y^* n = 14; *Atrx^R245C/y^* n = 13) and 56 weeks (*Atrx^+/y^* n = 13; *Atrx^R245C/y^* n = 13). On day 1, animals were placed in the testing chamber for 5 min and subjected to a 30 s tone (85 dB) ending in a 2 s 0.75 mA foot shock (programme: 120 baseline, 30 s tone ending in 2 s foot shock, 150 no stimuli). On day 2 (morning), contextual fear memory was assessed by returning animals to the same testing chamber for 5 min (programme: 300 s no stimuli); and (afternoon) cued fear memory was assessed by returning animals to a testing chamber that appeared different (by adding black triangular insert and white floor wiping with acetic acid) and playing them the same tone for 3 min (programme: 120 s baseline, 180 s tone). Behaviour was monitored and analysed using VideoFreeze^TM^ Fear Conditioning software (Med Associates Inc).

### Quantification and statistical analysis

#### Analysis of patient mutations

The list of unique ATR-X syndrome patient mutations and their frequency (number of cases) were obtained from Gibbons et al., 2008.

#### Quantification of western blots

Densitometry analysis was performed using ImageJ. Genotypes were compared using t-tests (unpaired, two-tailed).

#### Quantification of immunofluorescence

Cell counts were performed on 3–5 sections per animal and a minimum of 3 mice per genotype were used. Counts were expressed as a percentage of total DAPI cells unless stated otherwise. Coronal sections from control and mutant animals were first matched using age appropriate and specific brain landmarks located outside of the cortex. Following immunostaining, an identical sized box was oriented over the dorsomedial region of the telencephalon within which DAPI+ and marker+ cells were counted for all genotypes. For Purkinje cell counts, absolute numbers of calbindin+ cells were quantified across cerebellar lobes I to VII using Image J to quantify the distance of the Purkinje cell layer. Cell number was then presented as the average number of calbindin+ cells per 100 µm. The thickness of the corpus callosum was assessed rostrally (Bregma 0.86), medially (Bregma 0.5) and caudally (Bregma 0.26) by averaging 3 measurements per location. Genotypes were compared using t-tests (unpaired, two-tailed).

#### Analysis of alpha globin loci

Dot plots show sequence identity within the human and mouse alpha-globin loci and were generated using the Pustell DNA Matrix method in MacVector version 13.0.7 using the following parameters: Scoring matrix: DNA identity; Window size: 30; Hash value: 8; Minimum similarity: 60%.

#### Analysis of MRI data

After MRI, all images were corrected for geometric distortion (Nieman et al., 2018). Registration for image alignment was then performed using the Pydpiper toolkit (Friedel et al., 2014), generating a consensus average across all mice in the data set. Individual volumes were segmented by mapping onto a mouse brain atlas with 183 individual segmented structures (Dorr et al., 2008; Richards et al., 2011; Steadman et al., 2014; Ullmann et al., 2013) an automatically generated template approach (Chakravarty et al., 2013). Volumetric comparisons were then made using multiple t-tests as implemented in the R statistical computing package. Statistical significance was assessed after correction for the multiple comparisons using the false discovery rate (Benjamini and Hochberg procedure).

Cortical thickness was computed at each point on the outer surface of the cerebral cortex using the method described in (Lerch et al., 2008). Briefly, this method defines the inner and outer cortical surface as electrostatic potentials and solves for the length of field lines connecting the two surfaces.

Additional MRI data was obtained from Tamming et al., 2020 for comparison with *Atrx^fl/y^; CaMKIICre* (*CaMKII-cKO)* mice.

#### Sholl analysis

Cultured Hippocampal neurons were stained for MAP2 (1:1000, AB5622 Merck Millipore) and Tau-1 (1:1000 MAB3420, Merk millipore) to distinguish dendrites and axons, respectively and counterstained with Hoescht 33342 dye (Thermo Fisher Scientific). z-stack images were obtained using Zeiss AxioObserver Z1 and later processed using SNT plugins on FIJI to obtain a traced 2D binary representation of the dendrites for Sholl analysis using 10 µm steps and the number of crossings were counted until a radius of 150 µm, we also evaluate, total length and axonal length.

#### Statistical analysis of morphological and behavioural data

Growth curves were compared using repeated measures ANOVA. DEXA, Brain weights, tibialis anterior (TA) muscle weights and TA/body weights were compared using t-tests (unpaired, two-tailed).

X-ray images were measured using MicroDicom software. Body length was measured from nose to base of tail and genotypes were compared using a t-test (unpaired, two-tailed). Cranial index (%) = (cranial width/cranial length x 100). Zygomatic index = (zygomatic width/zygomatic length x 100), where zygomatic length is the mean of the left and right measurements. Kyphotic index = AB/CD, where AB is the length of a line drawn from posterior edge of C7 to the posterior edge of L6 and CD is the distance from line AB to the dorsal border of the vertebral body farthest from that line (drawn perpendicular to line AB)(Laws and Hoey, 2004). When data fit a normal distribution (cranial length, zygomatic length, zygomatic index, and kyphotic index), genotypes were compared using t-tests (unpaired, two-tailed). Otherwise (cranial index), genotypes were compared using KS tests.

The mean of three grip strength recordings (gram-force, GF) was normalised by body weight and genotypes were compared using t-tests (unpaired, two-tailed). Gait analysis parameters were exported from DigiGait Analysis 15 software and genotypes were compared using t-tests (if normally distributed) or KS tests. Stride length, analysed by t-tests (unpaired, two-tailed) and stance wide coefficient of variation (Cv), analysed by a KS test (unpaired, two-tailed) at 10 weeks and a t-test at 55 weeks are shown. Treadmill performance during trials (days 5-7) was compared for each day using Mann-Whitney tests. Feret’s diameter from TA muscle after chronic exercise was compared using repeated measures ANOVA.

Spontaneous locomotor activity in SHIRPA was compared using t-tests (unpaired, two-tailed). In the Open Field assay, distance travelled was assessed in four 5 min bins and genotypes were compared using repeated measures ANOVA. For each bin, genotypes were compared using t-tests (unpaired, two-tailed). In the Open Field assay, time spent in the central 40% of the arena was assessed in four 5 min bins and genotypes were compared using repeated measures ANOVA. In the Light/Dark test, time spent in the light half was assessed in three 5 min bins and genotypes were compared using repeated measures ANOVA. In the Elevated Zero Maze, time spent in the open quadrants was compared using a t-test (unpaired, two-tailed).

Baseline motor function was assessed by accelerating rotarod performance (latency to fall) during four trails on day 1. At 13 weeks, the data fit a normal distribution and so genotypes were compared using repeated measures ANOVA and improvement over the course of the day was assessed for each genotype using 1-way ANOVA. At 55 weeks, the data do not fit a normal distribution and so genotypes were compared for each trial using KS tests and improvement over the course of the day was assessed for each genotype using Friedman tests. Motor learning ability was assessed over the total three days. Daily performance for each animal was determined by the mean performance (latency to fall) in the four trials for each day. At 13 weeks, the data do not fit a normal distribution and so genotypes were compared for each day using KS tests and motor learning over the course of the experiment was assessed for each genotype using Friedman tests. At 55 weeks, the data fit a normal distribution and so genotypes were compared for each day using t-tests and motor learning over the course of the experiment was assessed for each genotype using 1-way ANOVA. Spatial learning was assessed in the Barnes maze by the time taken for the mouse to locate the escape hole, defined by its nose entering a zone around the hole. A maximum score of 180 s was assigned to animals that failed to locate the hole during the trial. Daily performance for each animal was determined by the mean time in the four trials for each day (days 2-5). Genotypes were compared on each day using KS tests. Social novelty preference was assessed by the amount of time the subject spends interacting with stranger mice (i.e. its nose is located in the sniffing zone around the cup containing the stranger mouse). Time spent sniffing stranger 1 vs 2 was assessed for each genotype using t-tests (paired, two-tailed). Spatial reference memory was assessed over 8 min in the Y-maze test. Alternation index = number of alternations/max alternations * 100. Genotypes were compared using t-tests (unpaired, two-tailed). Fear memory was determined by the time spending freezing minus the baseline the day after being subjected to foot shock, in response to being returned to the same chamber (contextual) or hearing the same tone (cued). In the contextual paradigm, time spent freezing was assessed in 1 min bins over the 5 min trial (minus baseline determined on day 1 prior to stimuli. Genotypes were compared using repeated measures ANOVA. In the cued paradigm, time spent freezing was assessed during the 3 min tone (minus baseline determined in the same trial prior to the tone). At 13 weeks data do not fit a normal distribution at genotypes were compared using a KS test. At 56 weeks data fit a normal distribution at genotypes were compared using a t-test (unpaired, two-tailed).

All statistical analysis of morphological and behavioural data was performed using GraphPad Prism 9. Significance thresholds: not significant (NS) P > 0.05; * P < 0.05; ** P < 0.01; *** P < 0.001; **** P < 0.0001.

